# Genome-wide DNA Methylation Patterns in *Daphnia magna* are Not Significantly Associated with Age

**DOI:** 10.1101/2024.06.27.600920

**Authors:** Ruoshui Liu, Marco Morselli, Lev Y. Yampolsky, Leonid Peshkin, Matteo Pellegrini

## Abstract

Studying DNA methylation in *Daphnia magna*, a model organism in ecological and evolutionary research, offers valuable insights into pharmaceutical toxicity and behavioral ethology. In this study, we characterized DNA methyltransferases and mapped DNA methylation across the *D. magna* genome using whole-genome bisulfite sequencing. Our analysis revealed a highly expressed, nonfunctional *de novo* methyltransferase (DNMT3.1) alongside lowly expressed functional *de novo* methyltransferase DNMT3.2 and maintenance methyltransferase DNMT1. *D. magna* displays overall low DNA methylation, targeting primarily CpG dinucleotides. Methylation is sparse at promoters but elevated in the first exons downstream of transcription start sites, with these exons showing hypermethylation relative to adjacent introns. In contrast to prior studies, we observed minimal age-related changes in DNA methylation patterns that were not sufficiently robust to build an accurate epigenetic clock. These findings expand our understanding of the epigenetic landscape in *D. magna*.

## INTRODUCTION

*Daphnia* are planktonic crustaceans within the Phyllopoda subclass and the Branchiopoda class (1). Among over 100 species within the *Daphnia* genus, *Daphnia magna* (*D. magna*) is widely recognized as a model organism in biological research, including studies on pharmaceutical toxicity, reproductive cycles, behavioral ethology, and phenotypic plasticity (2). DNA methylation, the addition of a methyl group on the 5’ carbon of cytosine, plays a role in transcriptional regulation and phenotypic expression (3). Previous studies have demonstrated that DNA methylation assessment aids in toxicity evaluation (4). Given the significance of *D. magna* as a model organism, examining its epigenomics landscape, particularly DNA methylation pattern and regulation, could be critical (5).

CpG methylation is closely associated with gene silencing and is crucial for development (6,7). In *Daphnia magna*, cytosine methylation levels are relatively low, yet specific regions, such as exons, exhibit higher methylation (3,6,8). Previous studies, however, have tended to average methylation across different genes for each exon, overlooking the nuanced methylation patterns of individual genes and exons. To address this gap, our study aims to examine both average methylation patterns and detailed variations at the exon and gene levels across different genes.

DNA methyltransferases (DNMTs) are a family of enzymes that epigenetically regulate gene expression by establishing and maintaining CpG methylation patterns (9). Maintenance methyltransferase DNMT1 replicates DNA methylation patterns to daughter cells during cell divisions, while the functional *de novo* methyltransferase DNMT 3 establishes methyl groups at unmethylated cytosine sites (10,11). Our research aims to explore the functional domains of *D. magna*’s DNMTs and their effect on genome-wide and gene-specific methylation patterns.

DNA methylation levels vary with age and influence the functional capability of organs in mammals, positioning DNA methylation-based biomarkers, also known as epigenetic clocks, as effective estimators of biological age (12–14). Epigenetic clocks facilitate understanding the impact of both endogenous (epigenetic drift, etc.) and exogenous stressors on biological aging by comparing epigenetic age to chronological age (12,14). In a previous study, an epigenetic clock for *D. magna* was built with 12 clock CpGs (6). However, this analysis was conducted using samples from only two age groups: 10-day and 50-day-old. Additionally, they incorporated *Daphnia* specimens from two different strains and included an interaction term for strain and age in their epigenetic clock. Both the large age gaps and an additional factor of strain potentially diminished the predictive accuracy of their regression model (6). Thus, further research is needed to characterize age-related changes in DNA methylation and determine the validity of an epigenetic clock for *D. magna*.

This study aimed to investigate the targeting and age association of DNA methylation in *D. magna*. We cultured 17 *D. magna* samples of various ages, performed whole genome bisulfite sequencing, and characterized the DNA methylation patterns at the genome-wide, gene, and exon levels. To better understand the methylation patterns, we analyzed functional domains and mRNA expression levels of DNA methyltransferases. Lastly, we examined age-related changes in methylation patterns and attempted to build an epigenetic clock.

## MATERIALS AND METHODS

### Sample collection

A heat-tolerant clone of D. magna, IL-MI-8, was originally acquired from the Ebert Lab at the University of Basel, Switzerland. This clone was sourced from a pond in Jerusalem, Israel, as part of the Ebert Lab stock collection. To ensure synchronized cohorts, neonates born within a two-day window were separated from their mothers. Sex was determined at 8- to 9-day-old animals. Female individuals were exclusively used in all experiments conducted in this study. All mothers received adequate nutrition and were cultured under identical conditions at 25°C within an incubator. All cultures, including mother cultures, neonate cultures, and tank cultures, were maintained in ADaM water (15). Animals were exposed to a light cycle consisting of 16 hours of light followed by 8 hours of darkness. Animals were fed a daily suspension of the green alga Scenedesmus obliquus at a concentration of 10^5 cells/ml (adjusted for population density, with one animal per 20 ml). Every sixth day, the water was replaced, and offspring were manually removed until animals were transferred to the culture platform. The operational protocols for the culture platform are akin to those described in previously published protocols (16).

### DNA extraction and WGBS library preparation

Approximately 30-50 ng of purified genomic DNA in 50 µl has been subject to sonication using the Bioruptor Pico (Diagenode) for 15 cycles (30 ON; 90 OFF). Fragmentation patterns have been checked with the D1000HS Assay (Agilent Technologies) on a 4200 TapeStation. The volume of the fragmented DNA has been reduced to 20 µl using a Vacufuge (Eppendorf), then subject to bisulfite conversion using the EZ DNA Methylation-Lightning Kit (Zymo Research). The libraries have been prepared using the Accel-NGS Methyl kit (Swift Biosciences - now xGen Methyl-Seq Library Prep - IDT) according to the manufacturer’s recommendations except for a major modification. Briefly, the denatured BS-converted gDNA was subject to the adaptase, extension, and ligation reaction. Following the ligation purification, the DNA underwent primer extension (98C for 1 minute; 70C for 2 minutes; 65C for 5 minutes; 4C hold) using oligos containing random UMI at the location of the i5 barcode. The extension using a UMI-containing primer allows the tagging of each molecule to remove PCR duplicates and correctly estimate DNA methylation levels. Following exonuclease I treatment and subsequent purification, the libraries were then amplified using a universal custom P5 primer and i7-barcoded P7 primers (initial denaturation: 98C for 30 seconds; 10 cycles of 98C for 10 seconds, 60C for 30 seconds, 68C for 60 seconds; final extension: 68C for 5 minutes; 4C hold). The resulting single-indexed libraries were then purified and quantified using the Qubit HS-dsDNA assay, and the quality was checked using the D1000-HS assay (Agilent - TapeStation 4200). The libraries were sequenced as 150 PE on the Illumina NovaSeq6000 platform.

### Bisulfite sequencing data processing

An alignment index incorporating a bisulfite-converted sequence for each reference strand was constructed from the *Daphnia magna* Xinb3 reference genome (BioProject ID: PRJNA624896, D. Ebert, personal communication) and the mitochondrial genome (17), using the *BSBolt Index* tool v1.6.0 (18). The FASTQ files were aligned using *BSBolt Align* (default options) (18). PCR duplicates were removed with *samtools markdup* v1.17 (option -r) (19). Sequence alignment quality was assessed using the CIGAR (compact idiosyncratic gapped alignment report) strings, and sequences with a total number of matches below 50 were excluded from further analysis using *samtools view* and *samtools index* (19). The number of mapped reads in each sample was quantified using *samtools flagstat* (19).

DNA methylation calling was performed with *BSBolt CallMethylation* (options: -IO) with the DNA alignment index as a reference, generating CGmap files (18,20). The CGmap files were subsequently filtered based on contig coverage and quality. Average coverage before and after filtering for cytosines across each chromosome was obtained using *CGmaptools mec* v0.1.2 (20). Global DNA methylation levels were analyzed with *CGmaptools mstat* (20), and differences in global methylation levels of CpG and non-CpG cytosines were assessed using the Wilcoxon test. DNA methylation levels of genes, exons, and introns were calculated using *CGmaptools mtr* (20). Metagene plots for genes and exons were generated using *CGmaptools bed2fragreg* and *CGmaptools mfg* (20). Pathway enrichment analysis was performed using the R package *clusterProfiler* (21). Genome browser snapshots were taken from Integrative Genomics Viewer (version IGV_2.18.2) (22).

The matrix of common CpG sites was generated using *BSBolt AggregateMatrix* function (options: -min-coverage 10 -min-sample 0.8 -CG) (18). To handle missing data, CpG sites absent in some samples were imputed using the *BSBolt Impute* feature. UMAP was employed to visualize the clustering of samples based on methylated CpG sites. This approach allowed us to examine whether *Daphnia* samples from the same age group exhibited consistent methylation patterns, with an expectation of tighter clustering within age groups in the UMAP plot (23).

The CpG sites were filtered to only include the ones with the top 20% of the variability. The epigenetic clock model was built with the Lasso function from the Python module *sklearn.linear_model* and Leave-One-Out Cross Validation (24). Differentially methylated regions (DMR) analysis was performed using the *Metilene* package v0.2-8 (options: -m 10 -d 0.001 -t 4 -f 1) (25).

Random permutation was used to assess the statistical significance of differences in methylation levels between age groups at each position within the meta-gene ridge plot. For each position, we calculated the differences in mean methylation levels between the two age groups. We conducted 100,000 random permutations and reassigned sample labels to form two new groups, ensuring that the size of these groups matched the age groups. In each permutation, we calculated the mean methylation difference between these newly assigned groups. p-values were calculated as the proportion of permuted differences as extreme as, or more extreme than, the observed differences, applying a two-tailed test. Significance at each site was defined as p-value < 0.05. This procedure was utilized across three comparisons: mature versus young, old versus young, and old versus mature.

### RNA sequencing data processing

Details of an age-specific RNAseq experiment are reported elsewhere (26). Briefly, RNA was extracted from somatic tissues (whole bodies) of *Daphnia magna* females from laboratory clone GB- EL75-69 (Basel University *Daphnia* Stock Collection, Switzerland) of different ages: young, reproducing (age 15-20 days), old, reproductively senescent, and old, reproductively rejuvenated (age both 130-175 days), in 4 replicates each. In all cases, *Daphnia* samples were sampled within 24 hours of molting and egg-laying and freshly laid eggs (if any) were removed from the brood chamber before homogenizing *Daphnia*. RNAs were extracted using Qiagen RNeasyPlus Mini kit and RNA library preparation was performed using NEBNext Ultra II DirecMonal RNA Library Prep Kit (NEB, Lynn, MA) following the manufacturer’s protocols. The libraries were sequenced with Illumina Novoseq 6000, S4 flow cell, PE100. Reads were mapped to *D. magna* Xinb3 reference transcriptome (BioProject ID: PRJNA624896; D. Ebert and P. Fields, personal communication), and genes with differential expression between young vs. old *Daphnia* were identified using DEseq2 (27). Statistical analysis of RNA expression levels between groups was performed using a two-sided t-test.

### DNA methyltransferase analysis

The functional analysis of DNMT3.1, DNMT3.2, DNMT1, and UHRF1 was conducted using InterProScan (28). Gene sequences were input into the tool to identify and characterize the functional domains within these methyltransferases. Nuclear localization signals were predicted using NLStradamus, a Hidden Markov Model (HMM) tool, to assess their potential nuclear targeting (29). The analysis was conducted using a 2-state HMM static model, and NLS sequences were identified based on a consensus between the Viterbi path and posterior probability estimates exceeding 70%.

## RESULTS

### *D. magna* has three DNA methyltransferases

Based on the homology with other organisms’ DNA methyltransferases (DNMTs), *D. magna* harbors two functional and one non-functional DNMT (**Table S1**) (9). It has two *de novo* DNMTs, *Dma*DNMT3.1 and *Dma*DNMT 3.2 (**Figure 1A**) (10).

**Figure 1.**
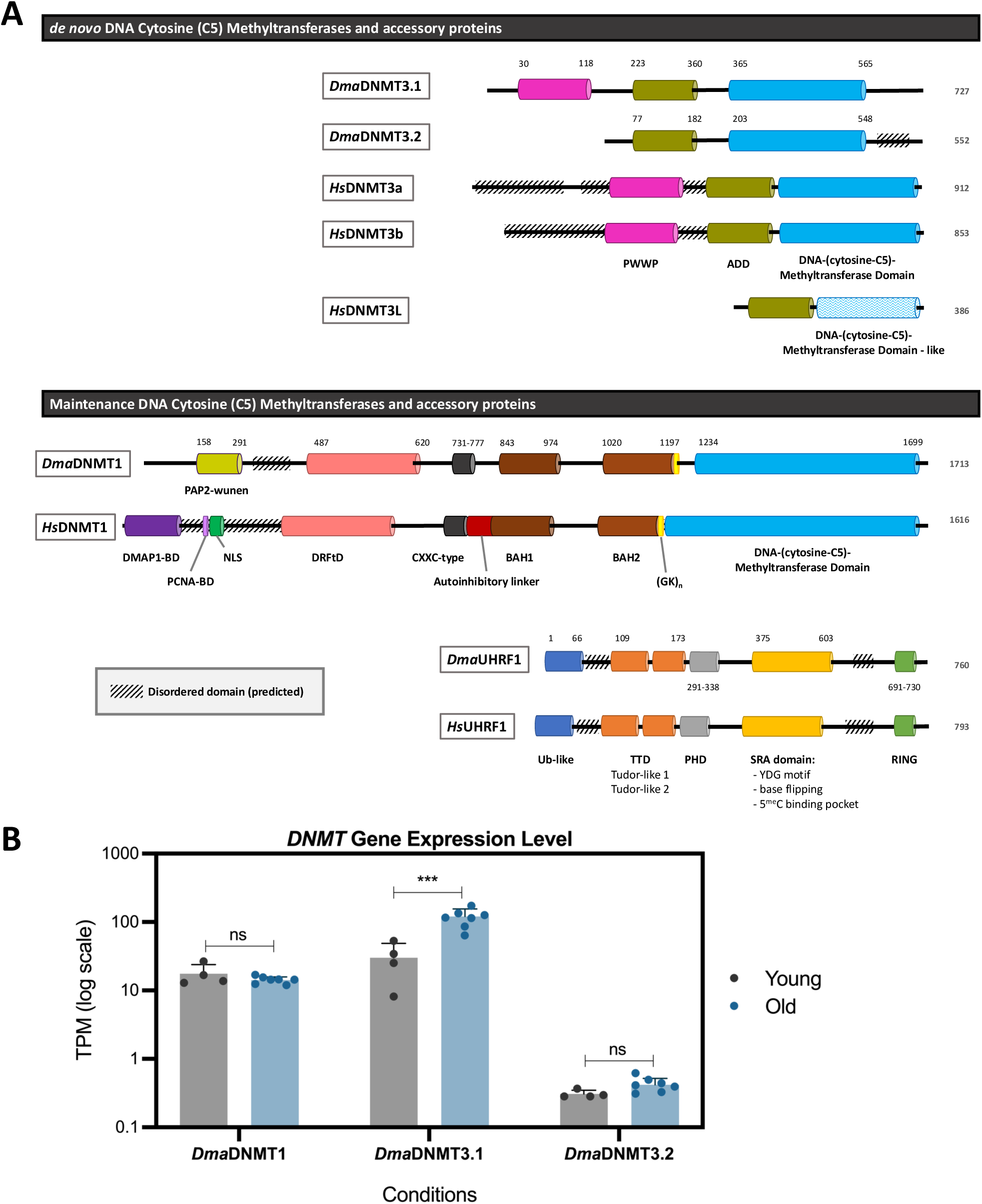
The comparison of DNA (C5) methyltransferases and accessory proteins of *D. magna* and *H. sapiens* and the gene expression levels under different conditions. **(A)** Domain structures of *de novo* and maintenance DNA methyltransferases and accessory proteins in *Daphnia magna* and *Homo sapiens* were predicted using InterProScan. Colored boxes represent the functional domains, with numbers indicating their respective positions along the protein sequence. Diagonal black-striped boxes indicate a predicted disordered consensus sequence. **(B)** Gene expression levels (TPM on log scale) of three DNMT genes (*Dma*DNMT1, *Dma*DNMT3.1, and *Dma*DNMT3.2) in young and old *D. magna* individuals. Bars represent the mean expression levels, and error bars denote standard deviation (SD). Statistical significance between conditions for each gene was assessed using a two-sided Student’s t-test, with significance indicated as follows: P < 0.001 (***) and nonsignificant (ns). Expression levels of *Dma*DNMT1 and *Dma*DNMT3.2 showed no significant differences between the groups, while *Dma*DNMT3.1 expression was significantly different in young individuals compared to the old group.

To better understand DmaDNMT and its relationship with the species’ methylation pattern, we used mammalian (human) DNMTs as a reference and compared the homologies between *Dma*DNMTs and *Hs*DNMTs. Mammalian DNMT3a/b contains a Pro-Trp-Trp-Pro (PWWP) domain, an ATRX- DNMT3A-DNMT3L (ADD) domain, and a DNA-(cytosine-C5)-Methyltransferase (MTase) domain with 6 motifs (I to VI). Non-functional *Dma*DNMT3.1 is missing 5 critical motifs in the MTase domain, and it is thought to exert similar functions to mammalian DNMT3L. Both *Dma*DNMT3.1 and *Hs*DNMT3L contain putatively functional ADD domains that do not bind H3K4me3. However, *Dma*DNMT3.1 owns the PWWP domain that is known to bind methylated lysines, specifically H3K36me2/3 in mammals (11).

Conversely, the functional *de novo* DNA methyltransferase, *Dma*DNMT3.2, contains all six DNA methyltransferase motifs and the ADD domain, suggesting that it can sense the methylation status of H3K4 and bind unmethylated H3K4 histones (10). However, it is missing the N-terminal PWWP domain present in the active mammalian DNMT3A and DNMT3B.

The other putatively active DNA methyltransferase in *D. magna* is the maintenance methyltransferase *Dma*DNMT1 (**Figure 1A**). Compared to the mammalian counterpart, *Hs*DNMT1, it is missing the N-terminal domains involved in the targeting of the replication foci, including the DNA methyltransferase 1-associated protein-binding domain (DMAP1-BD) and the proliferating cell nuclear antigen-binding domain (PCNA-BD). NLStradamus predicted *Dma*DNMT1’s nuclear localization signals (NLS) both upstream (position 403-434) and downstream (678–696) of the DRFtD domain, with the upstream NLS aligning with the conserved position in *Hs*DNMT1. Additionally, a phosphatase-specific domain (PAP2-wunen) at the N-terminal region is unique to *Dma*DNMT1.

To understand the activity of the three *Dma*DNMTs, we analyzed RNA sequencing data across two age groups: young and old (**Figure 1B**). Gene expression levels (TPM) varied significantly across the three *Dma*DNMTs by a two-sided t-test. The two functional enzymes, *Dma*DNMT1 and *Dma*DNMT3.2, exhibited consistently low expression levels with no significant differences between the age groups. By contrast, the non-functional *Dma*DNMT3.1 showed significantly higher expression in the old group compared to the young group. However, since *Dma*DNMT3.1 lacks the critical DNA methyltransferase motifs, its expression is unlikely to impact methylation levels.

### High-coverage *D. magna* Methylomes

To determine the DNA methylation profiles of *Daphnia magna*, we obtained two 45-day-old (mature-old) Xinb3 clone individuals, Dap_S1 and Dap_S2. Whole-genome bisulfite sequencing was performed as described in Materials and Methods, and subsequent sequence alignment to the *D. magna* genome and methylation calling was carried out. On average, approximately 79% of the reads mapped primarily to the genome, and this dropped to 56% after removing duplicates and 53% after filtering by CIGAR (compact idiosyncratic gapped alignment report) strings. The *Daphnia magna* genome comprises 56.6 million cytosine sites across both strands. Applying a minimum coverage threshold of 10 reads per site in the methylation calling step, we successfully identified 41.1 million cytosines, representing approximately 72.6% of the total cytosine sites in the genome. Although the average coverage across contigs was around 60, the distribution demonstrated large variations (**Figure S1A**).

The original genome assembly of *Daphnia magna* consisted of one mitochondrial genome (17) and 608 nuclear contigs (BioProject ID: PRJNA624896, D. Ebert, personal communication), many of which are short. Approximately 50% of these contigs are under 20,000 base pairs and 25% are below 10,000 base pairs in length. Some short contigs showed pronounced high coverage and hypermethylation at CpG sites, suggesting that they are outliers and possible contaminants. To improve the quality of methylome, alignment filtration was implemented, retaining only the CpN sites from the top-performing 97 nuclear contigs validated by genetic recombination map, Hi-C data, and optical mapping (BioProject ID: PRJNA624896, D. Ebert, personal communication). Following selective filtration, 38.7 millions of these 41.1 million covered cytosine sites (94.1%) were retained for each sample. The mean contig length was 1.3 million base pairs, with an average coverage of approximately 60x (**Figure S1B**). This process reduced variability across contigs and confirmed the reliability of the remaining methylcytosine sites.

Consequently, the cytosine sites chosen for our downstream analysis were situated on the top-performing contigs with a minimum coverage of 10x and an average coverage of 65x (**Table S2, S3**). These selected sites were included in the analysis even if they did not appear consistently across both samples.

Besides the two deeply sequenced *D. magna* samples (Dap_S1 and Dap_S2), an additional 15 *D. magna* samples denoted Dap_D1 to Dap_D16 (Dap_D7 excluded due to low quality) were bisulfite sequenced across different age groups: young (9 days), mature (22 to 27 days), and old (51 to 58 days). Similar approaches of alignment, methylation calling, and filtration were applied to the 15 samples. This approach yielded an average of 10.9 million cytosine sites per sample, each with a minimum coverage of 10x, for our downstream analysis (**Table S2, S3**).

### Mitochondrial genome methylation as a negative control for non-conversion artifacts

Bisulfite treatment converts unmethylated cytosines to uracils, while leaving the methylated cytosines unchanged, allowing methylation to be accurately detected through sequencing. However, bisulfite conversion is not 100% effective, and a small fraction (typically around 1%) of unmethylated cytosines fail to deaminate and are mistakenly identified as methylated (30).

To distinguish between methylcytosines and unconverted cytosines we considered several criteria. If cytosine dinucleotides are methylated, we expect the methylation patterns at individual sites to be consistent across samples. To investigate this, we conducted correlation analyses between the Dap_S1 and Dap_S2 samples across different cytosine contexts. We observed a high correlation (r = 0.949) for CpG sites between the two samples, indicating reproducible methylation patterns. We also expect that methylated CpG dinucleotides are present on both strands, which is often referred to as symmetric methylation. In Dap_S1, we identified 1,100,329 pairs of CpG sites with non-zero methylation on both strands, of which 1,040,273 pairs (94.5%) had a difference in methylation of less than 0.05. Similarly, in Dap_S2, 1,204,966 CpG pairs had non-zero methylation, with 1,129,724 pairs (93.8%) showing a difference below 0.05. This high degree of symmetry between strands further supports the reliability of CpG methylation signals (31). By contrast, non-CpG contexts exhibited much lower correlations, with CpA at r = 0.338, CpC at r = 0.222, and CpT at r = 0.249. The low correlation levels for non-CpG sites suggest that these are likely artifacts of incomplete bisulfite conversion rather than true methylation.

Moreover, to differentiate between bisulfite non-conversion and true cytosine methylation, we used the mitochondrial genome as a negative control, as previous studies have shown that mitochondrial DNA is generally unmethylated (32) and have utilized this approach to assess true methylation level (33). Cytosine dinucleotides in the mitochondrial genome exhibited median coverage levels exceeding 100x (**Figure S2**), and this high coverage allows us to use the mitochondrial genome as a negative control. Dap_S1 and Dap_S2 displayed consistent methylation patterns across all cytosine contexts (**Figure S3**) with median methylation levels ranging from 0.2% to 0.5%. Non-CpG sites had third-quartile methylation levels between 1.0% and 1.2%, while CpG sites exhibited a slightly higher third quartile of 1.6%. To estimate the non-conversion rate in the nuclear genome, we compared the methylation levels of individual nuclear cytosine sites with those in the mitochondrial genome. The average methylation levels in the mitochondrial genome were comparable to or even higher than those observed in the nuclear genome (**Figure S4**). Moreover, non-CpG sites in the mitochondrial genome showed a higher percentage of methylation levels exceeding 20% compared to nuclear sites. However, the presence of nuclear mitochondrial sequences (nuMT) (34) in *D. magna*—mitochondrial DNA fragments inserted into the nuclear genome—may explain some of these effects (35). These nuMT sequences are fragments of mitochondrial DNA and thus contribute ambiguity to interpreting the data and potentially skewing nuclear methylation measurements by retaining mitochondrial methylation patterns or contributing additional unmethylated cytosines, depending on their conversion rates. To estimate the proportion of nuMTs, we re- mapped reads originally aligned to the mitochondrial genome onto the nuclear genome. We found that 26.1% of mitochondrial-aligned reads in Dap_S1 and 22.7% in Dap_S2 mapped to the nuclear genome with a normalized Alignment Score greater than 90%. This score was calculated by dividing the alignment score from BSBolt Align by the read length. This substantial fraction suggests that a portion of what appears to be mitochondrial methylation may instead originate from nuMTs.

In contrast to non-CpG sites, CpG sites in the nuclear genome displayed higher methylation levels than those in the mitochondrial genome, suggesting that CpG sites in the nuclear genome are methylated rather than artifacts of non-conversion.

### *D. magna* DNA CpG sites are methylated at low levels

To elucidate methylation at a global level, we computed the average methylation levels of the 97 contigs. CpT and CpC sites exhibited an average methylation level of approximately 1.25%, while CpA methylation was slightly lower with 1.1% methylation (**Figure S5**). By contrast, CpG sites displayed higher average methylation levels, around 2.05% across contigs, whose distribution of methylation was significantly different from that of the non-CpG sites (p-value < 0.0001 in Wilcox test).

We further assessed the methylation levels at individual CpN sites and categorized corresponding fractions of methylation levels (**Figure 2**). We found that the two samples had comparable methylation levels, and the methylation levels of CpA, CpT, and CpC sites were distributed similarly. Around 40% of the non-CpG sites had a non-zero methylation level below 0.05, while around 55% had no methylation. Approximately 5% of the non-CpG sites had a methylation level above 0.05 and less than 0.2% above 0.20. By contrast, CpG sites are significantly hypermethylated compared to the three non-CpG cytosine contexts in both samples (p-value < 0.0001 in the Wilcox test). Approximately 6% of the CpG sites had a methylation level above 0.05 and around 1% of the sites above 0.20. Specifically, 0.5% of the CpG sites have a methylation level above 0.80. This higher methylation frequency in CpG sites could be attributed to the specificity of DNMTs, which primarily target CpG dinucleotides over non-CpG sites (36). This pattern emphasizes the established role of CpG methylation in invertebrate transcriptional regulation (37), further distinguishing it from non-CpG methylation contexts.

**Figure 2.**
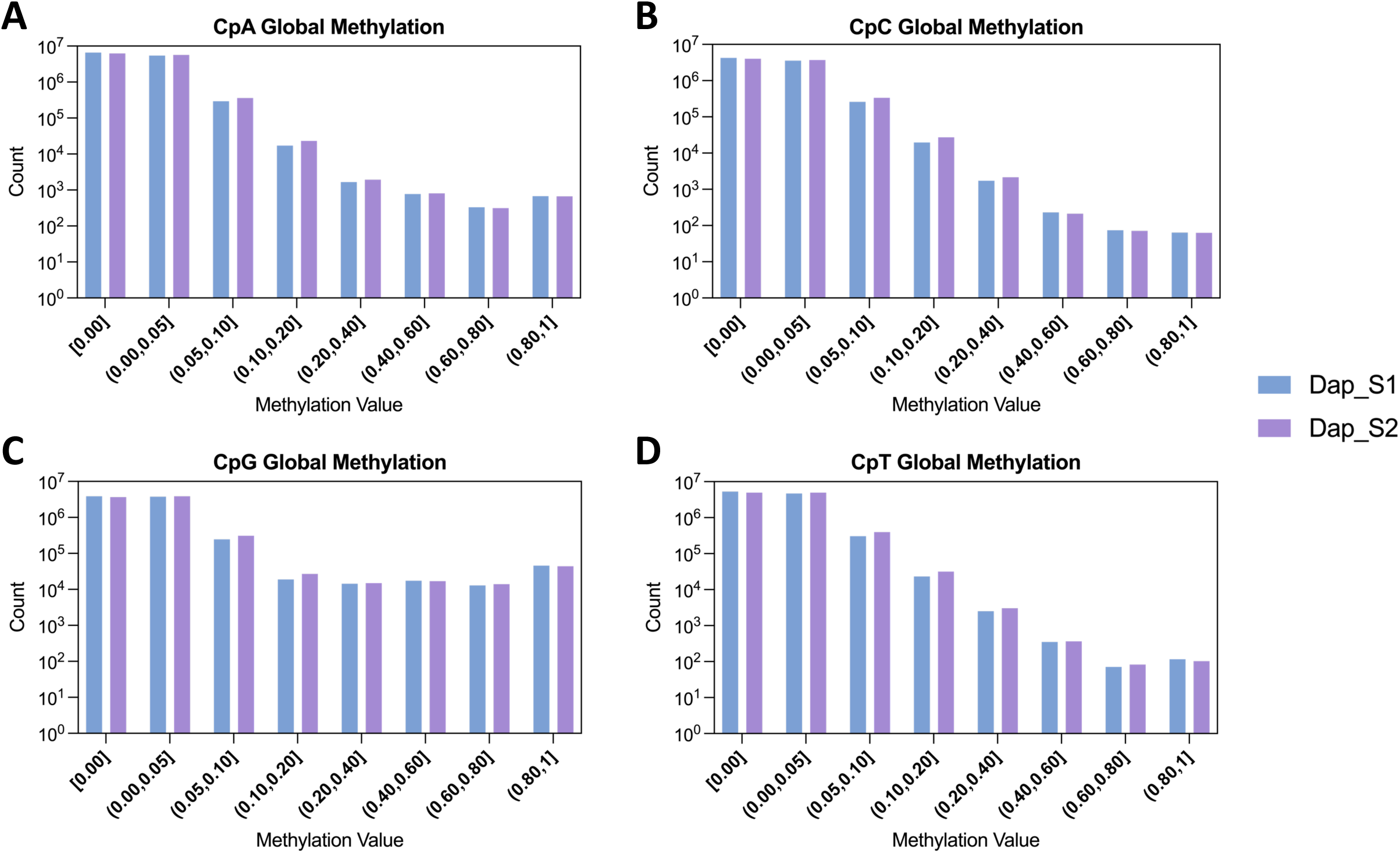
Global DNA methylation levels at CpN sites of two deeply sequenced samples. The count distribution of methylation levels for individual CpN sites is shown. **(A)** CpA, **(B)** CpC, and **(D)** CpT sites exhibit comparably low methylation levels. **(C)** CpG sites display a significantly higher degree of methylation overall.

In addition to examining the overall methylation trends, we analyzed the methylation profiles within specific groups of sequences, including DNA transposons, retrotransposons, LTR retrotransposons, and non-LTR elements. While some short interspersed nuclear elements (e.g., SINE_U, SINE_B2, SINE_Alu) showed localized methylation upstream or downstream of the sequences, most repetitive elements exhibited constant low methylation levels.

### *D. magna* DNA CpG sites are hypomethylated upstream of TSS

We next examined methylation patterns around genes, focusing on the range of 3 kilobases upstream of the transcription start site (TSS) to 3 kilobases downstream of the transcription end site (TES). Both samples, Dap_S1 and Dap_S2, showed consistent trends, highlighting the reproducibility of these patterns. Non-CpG sites maintained a uniform methylation level of approximately 1.5% across the gene and proximal regions (**Figure 3A**, **Figure 3B**, **Figure 3D**), suggesting non-CpG methylation is mainly an artifact and represents background noise. In contrast, CpG methylation displayed greater variability at different positions along genes. Specifically, CpG methylation levels were lowest a few hundred bases upstream of the TSS (1.5% methylated) and sharply increased downstream of the TSS, peaking at nearly 4.5% (**Figure 3C**). Subsequently, within the gene, CpG methylation levels gradually decreased, reaching 1.5%, underscoring the enrichment on the 5’ end of the genes.

**Figure 3.**
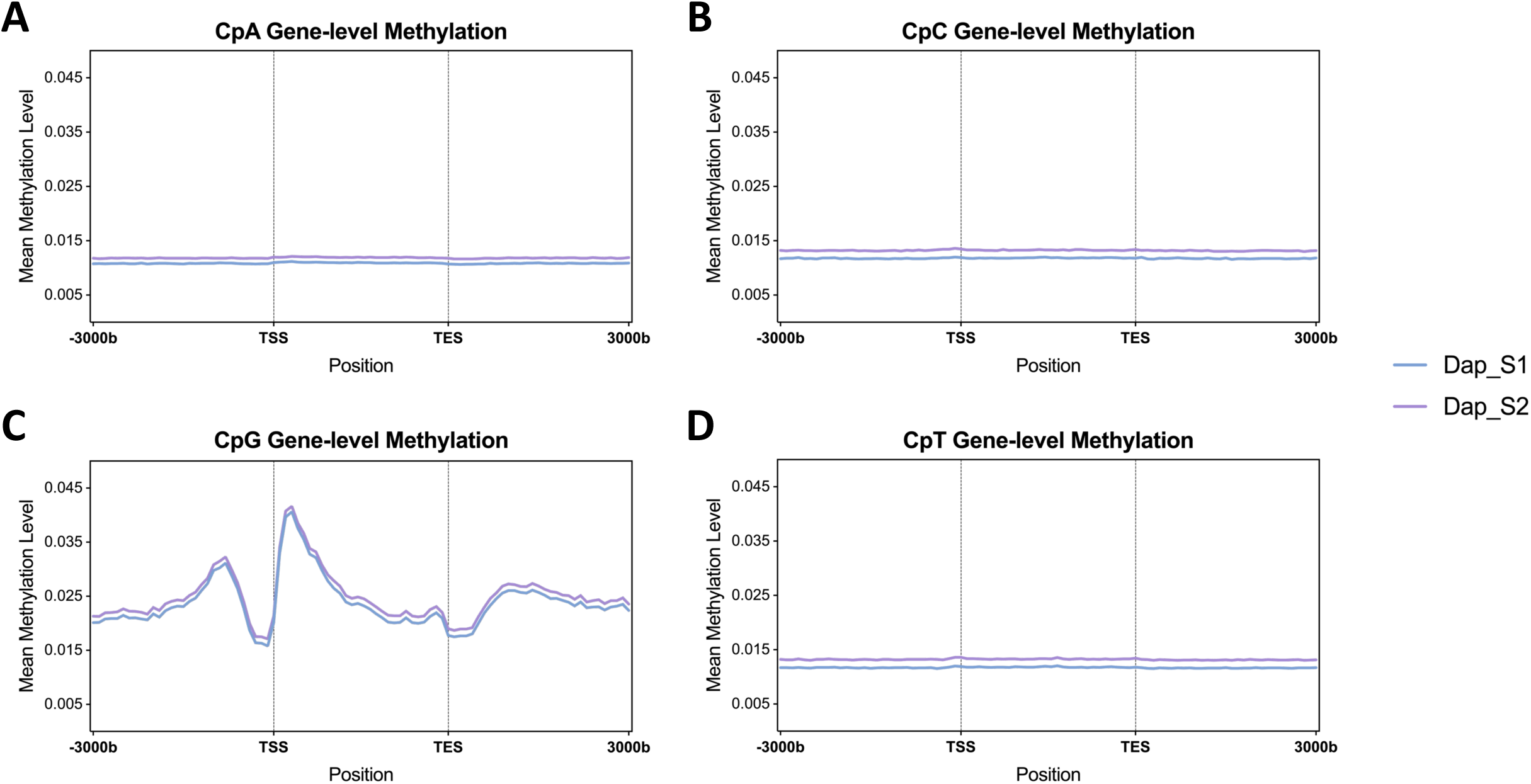
Gene-level DNA CpN methylation levels in two deeply sequenced samples, spanning 3 kilobases (kb) upstream transcription start sites (TSS) and 3 kb downstream of transcription end sites (TES). Each line depicts the average gene methylation value for **(A)** CpA, **(B)** CpC, **(C)** CpG, and **(D)** CpT for a sample. Non-CpG sites exhibited uniformly low methylation levels, whereas CpG sites manifest a distinctive pattern. These sites have lower methylation levels at TSS compared to regions upstream and downstream.

To minimize positional artifacts from normalizing genes to a uniform distance and to provide more insight into dynamic CpG methylation within genes, we grouped genes into deciles based on their mean methylation levels (e.g., top 10%, 10-20%) and calculated methylation levels centered at the TSS and TES. Dap_S1 (**Figure 4**) and Dap_S2 (**Figure S6**) displayed high methylation in the top 20% of genes (**Figure S7**) and minimal methylation in the lower 50% of genes (**Figure S8**). Around the TSS, the top 20% of genes showed markedly higher methylation levels than the rest, with methylation at the 5’ end exceeding 30% in the top 10% and nearly reaching 20% in the 10-20% group. At the same time, methylation in the top 10% of genes peaked at 10% upstream of the TES.

**Figure 4.**
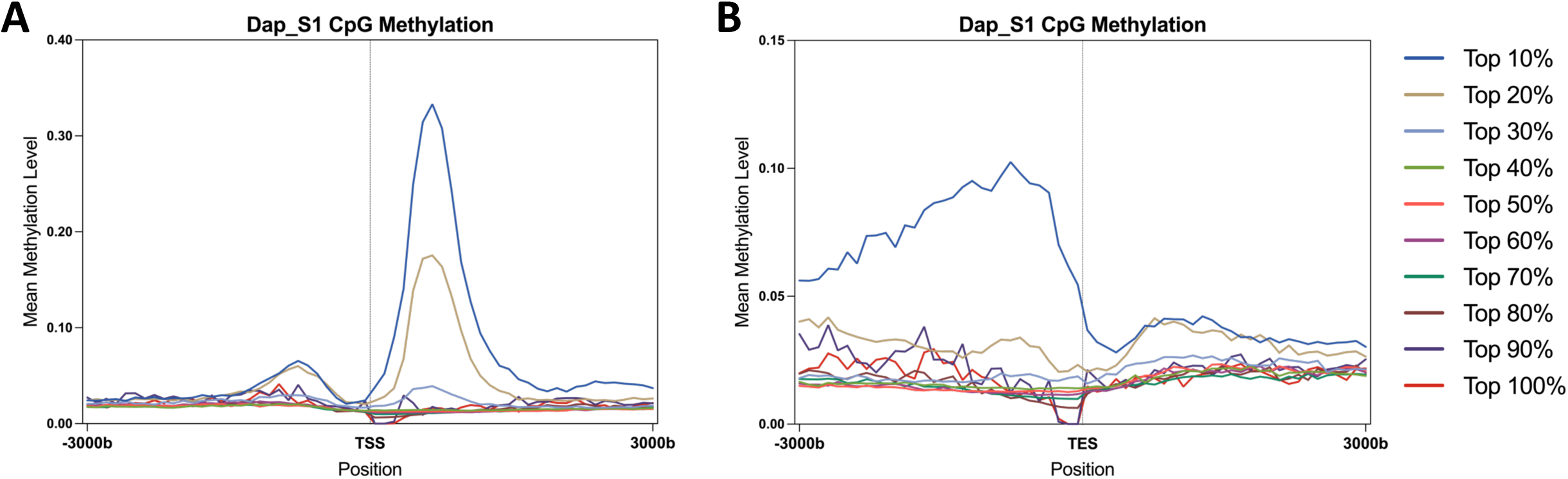
Gene-level CpG Methylation Patterns Centered around TSS and TES for Dap_S1, spanning 3 kilobases (kb) upstream and downstream of transcription start sites (TSS) and transcription end sites (TES). Genes were divided into 10 groups based on their mean methylation levels. Each line represents the average methylation level within each group, centered around **(A)** the transcription start site (TSS) and **(B)** the transcription end site (TES). For example, Top 20% in legend represents the top 10-20% group.

We explored the functions of the highly methylated genes and unmethylated genes using pathway enrichment analysis. Human homologs of *D. magna* genes within the top 10% methylation category across both samples were significantly enriched for fundamental cellular processes in the Gene Ontology Biology Process (GOBP) database, including ribonucleoprotein complex biogenesis, cytoplasmic translation, mitochondrial gene expression, respiratory electron transport chain, methylation, and DNA replication. Unmethylated genes showed no enrichment for any specific pathway.

### *D. magna* Exon CpG sites are more methylated than neighboring intron regions

To further explore the landscape of genomic methylation, we analyzed differential methylation patterns between exons and introns. Our findings reveal that exons contain a higher density of methylated CpG sites (**Figure S9**). On average, exons showed a methylation rate of approximately 0.17%, whereas nearby intronic regions were significantly less methylated, averaging 0.07% (p < 0.001 in the Wilcox test). Among the exons with a mean methylation rate greater than zero, the average methylation rate is 8.7%, significantly higher than the 4.9% observed in methylated introns (p < 0.001 in Wilcoxon test).

We also analyzed methylation levels across exons in genes with at least five exons to understand how exon position affects methylation. We observed that methylation varied significantly with exon positions (**Figure 5A**). Specifically, Exon 1 shows a gradual increase in CpG methylation, peaking near its 3’ end, followed by a sharp drop at the transition to the adjacent intron. This pattern of exon hypermethylation and intron hypomethylation is primarily driven by the top 10% of methylated genes (**Figure 5B**). Methylation levels continued to rise in Exon 2, reaching a peak of nearly 8% in Exon 3 before decreasing (**Figure 5A**).

**Figure 5.**
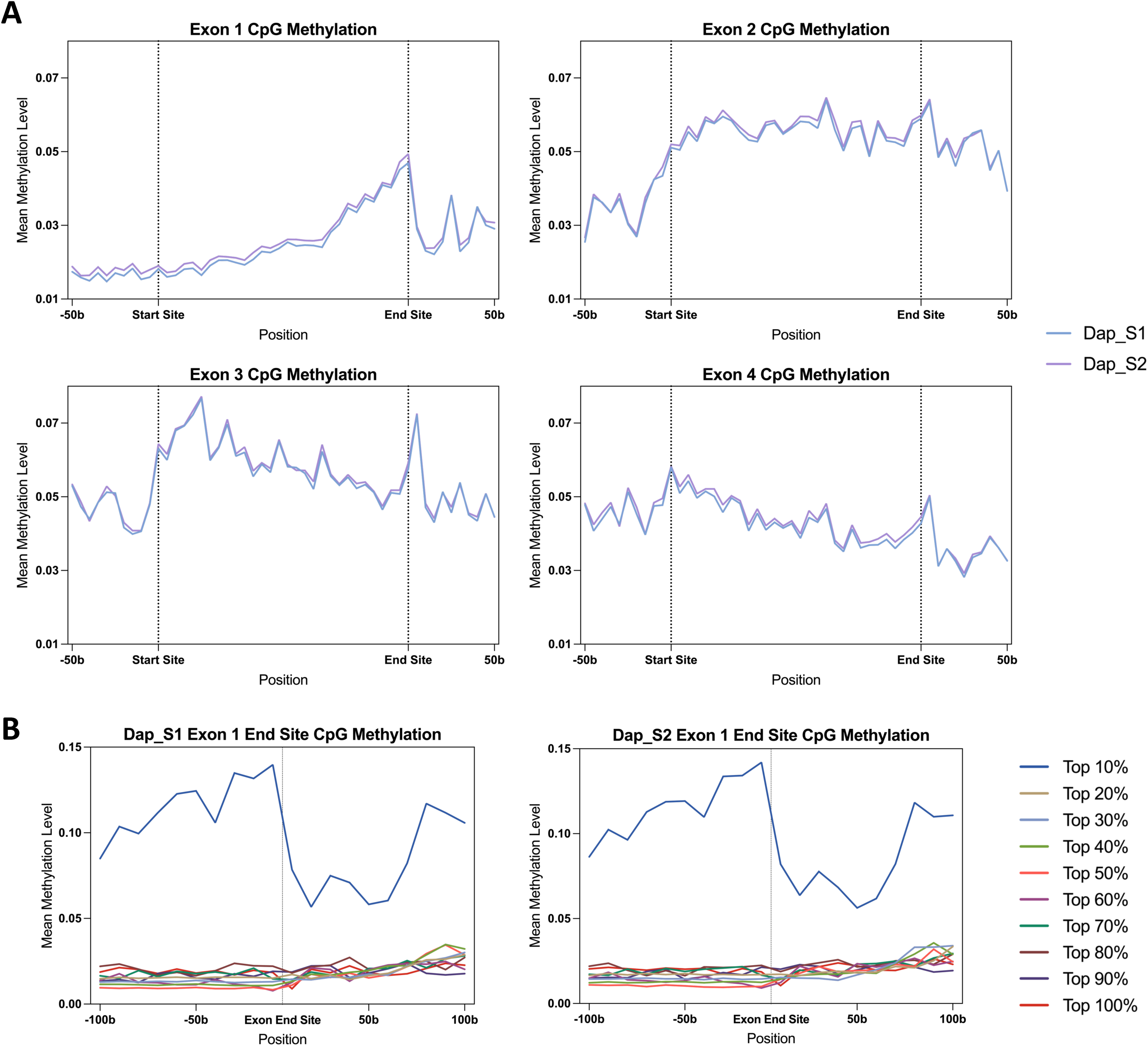
Exon-level CpG Methylation Patterns in two deeply sequenced samples. **(A)** CpG methylation levels of Exon 1 to Exon 4 from genes with more than 5 exons are demonstrated. Exon 1 shows a gradual increase in CpG methylation, peaking near its 3’ end. Methylation levels continued to rise in Exon 2, reaching a peak of nearly 8% in Exon 3 before decreasing. **(B)** Genes were divided into 10 groups based on their mean methylation levels in Exon 1. Each line represents the average methylation level within each group, centered around Exon 1 end site, with 100b up- and downstream of each region. In top 10% genes, there is a sharp drop in methylation transitioning from Exon 1 to the adjacent intron.

### *D. magna*’s global and local CpG methylation patterns are weakly associated with age

To determine how the methylation pattern changes with age, we first conducted a global analysis of CpG methylation levels across three age groups: young, mature, and old. The average methylation levels were 3.426% in the young group, 3.386% in the mature group, and 3.311% in the old group, showing a slight decrease with age; however, none of these differences reached statistical significance (**Figure 6A**). We also aggregated methylation values for CpG sites present in at least 80% of samples, imputed missing values, and constructed a CpG matrix. Visualization of this matrix using UMAP did not reveal clear clustering by age group, indicating a general absence of global age-related methylation patterns (**Figure 6B**). We further filtered the CpG matrix to retain CpG sites in genes with at least 10% methylation (562 genes in total) in at least 80% of the 17 samples. However, UMAP visualization of the filtered matrix of the filtered matrix also showed no age-related methylation patterns (**Figure S10**). Furthermore, to identify specific CpG sites that could serve as indicators of age, we trained an epigenetic clock through Lasso Leave-One-Out cross-validation regression (**Figure S11**). However, the correlation between predicted and actual ages was not significant, indicating that age-associated changes in the methylation of CpG sites were not robust enough to build an accurate epigenetic clock.

**Figure 6.**
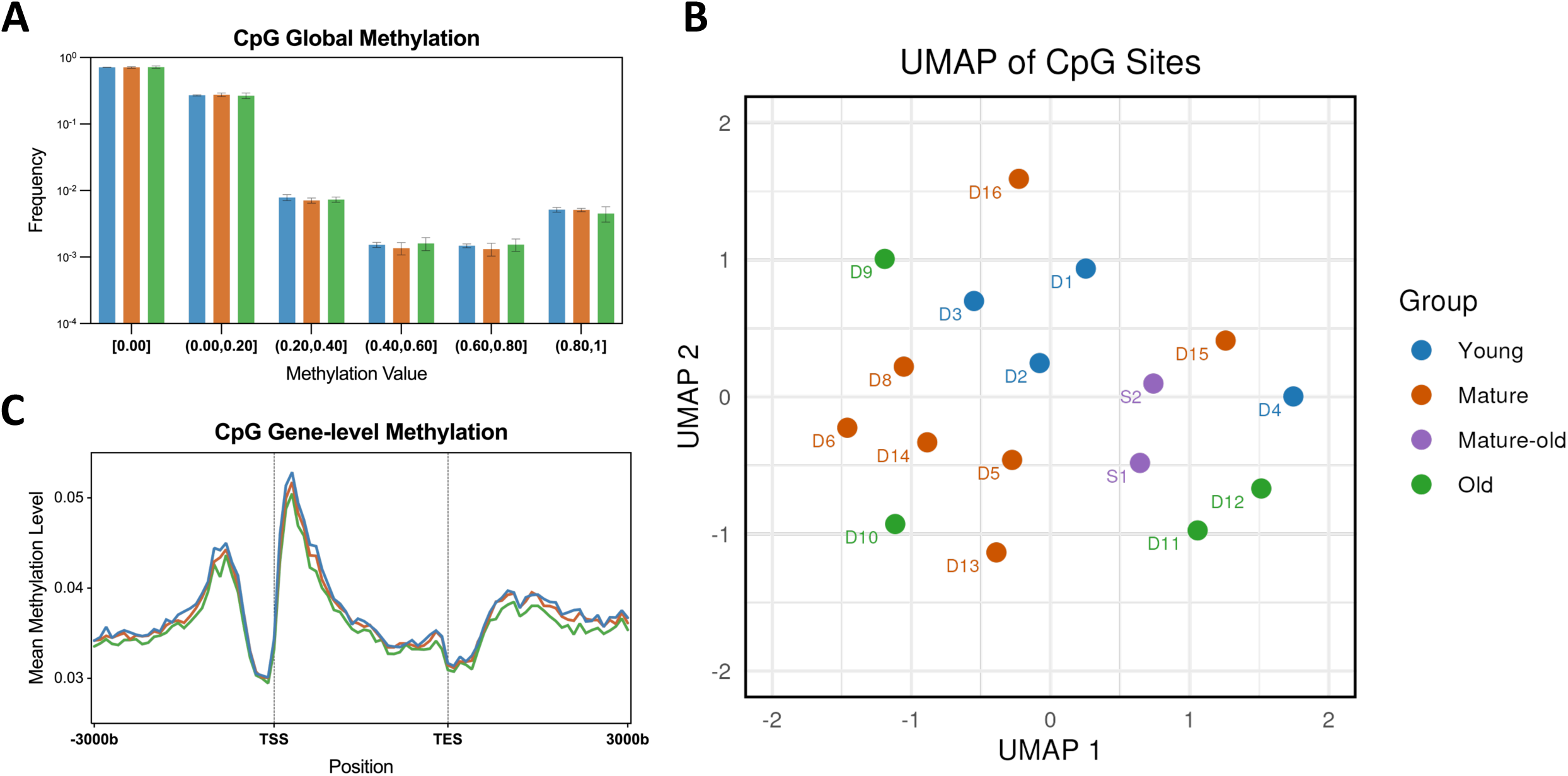
CpG methylation patterns across age groups demonstrate weak relation to age. **(A)** Methylation levels of individual CpG sites in each group are presented as the frequency distribution. **(B)** The UMAP visualization of matrix with CpG sites present in at least 80% of the 17 samples does not reveal any specific age-related clustering. **(C)** Meta-gene plot across age groups demonstrates a similar pattern.

Next, we explored age-related changes in CpG methylation at the gene level. While meta-gene methylation analysis demonstrated a consistent pattern across age groups (**Figure 6C**), closer examination of the differences between age groups revealed a weak trend of decreasing methylation with advancing age (**Figure S12**). However, random permutation tests at each position showed no statistically significant differences across all sites for the three pairwise age group comparisons. We also performed differentially methylated regions (DMR) analysis and linear regression analyses at the individual gene level, using the mean CpG methylation level as the dependent variable and age as the predictor. Neither any regions nor genes showed significant associations with age, indicating a lack of robust gene-level methylation changes.

## DISCUSSION

In this study, our primary focus was elucidating the genome-wide and gene-level methylation patterns in *Daphnia magna* and analyzing the age-associated variations in methylation levels. One key area of interest was the DNA methyltransferases in *D. magna*. Maintenance methyltransferase *Dma*DNMT1 and *de novo* methyltransferase *Dma*DNMT3.2 were both functional but expressed at low levels. In contrast, the non-functional DNMT3.1 exhibited higher expression levels and a potential association with aging, suggesting functions beyond DNA methylation. For example, DNMT3.1 has been shown to regulate growth and reproduction under starvation conditions (10).

To explore the evolutionary relationships of invertebrate DNA methyltransferases, we compared *Dma*DNMTs with DNMTs from other invertebrates. Within the phylum Arthropoda, species in the subphylum Crustacea, including *Daphnia pulex*, *Procambarus virginalis* (marbled crayfish), *Parhyale hawaiensis*, *Penaeus vannamei*, and *Hyalella azteca*, retain at least one DNMT3. In contrast, several crustaceans, such as *Armadillidium vulgare*, *Calanus finmarchicus*, *Eurytemora affinis*, *Lepeophtheirus salmonis*, and *Tigriopus californicus*, have lost DNMT3 (38,39). Expanding to the Hexapoda subphylum, which shares a common arthropod ancestor with crustaceans, DNMT3 is largely conserved in bees and ants, with *Solenopsis invicta* (red imported fire ant) and *Apis mellifera* (honey bee) possessing one DNMT3 (39–41). However, DNMT3 appears to be absent in most wasps, flies, butterflies, and moths, including *Polistes dominula*, *Drosophila melanogaster*, *Bombyx mori* (silkworm moth), and *Danaus plexippus* (monarch butterfly) (39,42,43). Moving further from Arthropoda, DNMT3 is also present in other invertebrate phyla. In Mollusca, *Crassostrea gigas* (oyster) retains functional DNMT3 (44). Similarly, in Annelida, *Capitella teleta* and *Ophryotrocha fusiformis* possess DNMT3, whereas *Dinophilus gyrociliatus* lacks it (45). The observed diversity in the presence and functionality of DNMT3 among invertebrates reflects adaptive modifications to their unique genomic and environmental demands, with *D. magna* representing one variation of this evolutionary landscape.

Most invertebrates possess a maintenance methyltransferase DNMT1 with a functional MTase protein domain. However, certain species, including *D. melanogaster*, *Aedes aegypti*, *Anopheles gambiae*, and *Caenorhabditis elegans* lack DNMT1 altogether (11,38–45). As observed in *D. magna*, the N-terminal DMAP-1 binding domain has been lost in many invertebrates, including arthropods (*A. mellifera*), annelids (*C. teleta* and *O. fusiformis*), ctenophores *(Pleurobrachia bachei*), brachiopods (*Lingula anatina*) (45–47). This supports previous findings that the DMAP-1 binding domain is conserved in chordates but largely absent in invertebrates (48).

We cultured 17 *D. magna* samples of different ages and conducted whole-genome bisulfite sequencing, achieving a high-coverage methylome with an approximate coverage of 60x. This allows us to analyze the methylation patterns across genome, gene, and exon levels. Globally, *D. magna* exhibited an average CpG methylation level of 2%. In comparison, mammals such as mice have an average CpG methylation level of around 80% (49). One possible explanation for this low methylation level is the reduced expression of *Dma*DNMT1 and the absence of the DMAP-1 binding domain. The DMAP-1 binding domain facilitates DNMT1’s interaction with DMAP-1, a co-repressor that aids its methylation maintenance activity (50). However, the correlation between the absence of DMAP-1 binding domain and reduced methylation appears weak and does not hold consistently across invertebrates. For example, the annelids *C. teleta* and *O. fusiformis* also lack no DMAP-1 binding domain, yet more than 17% of their CpG sites remain methylated (45). Similarly, *A. mellifera* lacks this domain but maintains global CpG methylation levels above 40% (51). These discrepancies suggest the factors contributing to *D. magna*’s low methylation levels remain unknown.

*D. magna* has an average gene body methylation of around 3%, slightly higher than the global methylation level. Gene body methylation refers to the addition of methyl groups, typically at CpG sites, within the coding regions of genes. It is often associated with actively transcribed genes, where it may help regulate splicing, reduce spurious transcription, and maintain genome stability (52). Gene body methylation is widely observed in animals, plants, and some fungi (33). The mosaic DNA methylation with a high level in gene bodies and lower level in intergenic regions was observed in many other invertebrates, including sea urchin, lancelet, honey bee *A. mellifera*, green peach aphid *Myzus persicae*, and *O. fusiformis and C. teleta* (45,53–55). However, *D. magna*’s gene body methylation is lower than the other invertebrates, which may be attributed to the absence of a functional PWWP domain in DNMT3.2. Since the PWWP domain binds to H3K36me2 and H3K36me3, markers typically found in transcribed regions, its absence could lead to a reduced recognition of H3K36-methylated gene bodies by the active DNMT3.2, resulting in low methylation levels (56–58). Supporting this, H3K36me3 enrichment in gene bodies has been observed in other invertebrates, including *Nematostella vectensis*, *Ciona intestinalis*, *A. mellifera*, and *Bombyx mori* (59).

In examining the methylation patterns across gene bodies and surrounding regions, we observed that the genes are hypomethylated upstream of the TSS and hypermethylated downstream of the TSS, with methylation levels decreasing along the genes. This pattern aligns with findings in the annelids *O. fusiformis* and *C. teleta*, where a similar trend of hypomethylation at TSS has been reported (45). The hypomethylation at the TSS likely results from a mechanism similar to that described in mammals, where the ADD domain of active DNMT3 selectively binds to unmethylated H3K4 regions (60). As mammalian promoters are typically rich in methylated H3K4, this prevents DNMT3 from binding and consequently blocks DNA methylation in mammalian systems (61). A comparable mechanism may operate in *Daphnia magna*, as previous research has shown a high enrichment of H3K4me3 upstream of the actively transcribed *Actin* gene (9). Furthermore, a strong correlation between gene expression and H3K4me3 levels at promoter regions in *Daphnia pulex* further supports this regulatory mechanism in *D. magna* (62).

We observed significantly higher methylation levels in exons compared to introns, which was also reported in *O. fusiformis*, *C. gigas*, and *A. mellifera* (44,45,49). This differential methylation is likely influenced by nucleosome positioning, as DNA methyltransferases preferentially target nucleosome- bound DNA (63). Consequently, the enrichment of nucleosomes on exons leads to higher methylation in these regions, suggesting a role for DNA methylation in exon definition and alternative splicing regulation (64,65).

Our analysis of *Daphnia magna* samples revealed minor, insignificant decreases in global and gene-specific methylation levels with age as well as no significant age-related clustering. This contrasts with vertebrates, where DNA methylation patterns are highly dynamic and exhibit notable changes during aging (66). Thus, the relatively constant DNA methylation level of *D. magna* across age underlines the possibility DNA methylation is less coupled to developmental changes in gene expression than in vertebrates, possibly due to the different structure of the DNA methyltransferases. At the same time, our attempt to build an epigenetic clock failed for *D. magna*. This failure is likely due to the low overall CpG methylation levels compared to mammals and the small magnitude of methylation changes with age, making it difficult to identify age-associated patterns necessary for clock construction. Indeed, only a few epigenetics clocks have been reported for invertebrates. Recently, an epigenetic clock on *Nasonia vitripennis*, the jewel wasp, using 19 age-predictive CpG sites was found to have a Spearman’s P of 0.94 for the correlation between predicted and actual age (67). However, the fact that the clock CpG sites were pre-selected for their age association indicates possible overfitting to their samples. Thus, we still lack robust evidence that epigenetic clocks can be generated for invertebrates.

Although no epigenetic clocks were established, we observed a slight decline in CpG site methylation levels with aging in *D. magna*. This pattern of epigenetic erosion is pronounced in invertebrates during development (68,69). For example, in *A. mellifera* (honey bee), exon methylation decreases from sperm and embryo to drone and worker larvae (70). Similarly, annelids such as *O. fusiformis* and *C. teleta* experience global methylation loss from embryonic to adult ages (45), while deuterostomes like sea urchin and lancelet display lower methylation levels in adulthood compared to earlier developmental stages (53). This feature of global hypomethylation has been reported as a consequence of heterochromatin loss, a hallmark of aging across diverse eukaryotes (71).

In this study, our primary emphasis was on the *D. magna* Xinb3 clone. The methylation profiles we elucidated may be slightly altered in other *D. magna* strains. Future research should explore the methylation patterns in additional clones to achieve a more comprehensive understanding. Given the limited correlation between DNA methylation and chronological age, our future attention will shift towards the influence of environmental factors, including temperature and water composition, on methylation dynamics.

## Declarations

LP was supported by the quadraScope Ventures fund and McKenzie Family Charitable Trust.

## Disclosure of interest

The authors declare that the research was conducted in the absence of any commercial or financial relationships that could be construed as potential conflicts of interest.

## Data availability of the Statement

Bisulfite sequencing data and CGmatrix generated in this study have been deposited within the Gene Expression Omnibus (GEO) repository, accession number GSE267859.

## Notes

### Competing Interest Statement

The authors have declared no competing interest.

### Summary of Updates

Figure 5 and 6 revised; results and discussion section updated; supplemental files updated

