## Supplemental Files for "Genome-wide DNA Methylation Patterns in *Daphnia magna* are Not Significantly Associated with Age"

### Supplementary Tables

**Table S1.** ***D. magna*** **DNA methyltransferases and accessory protein sequences.**

| **Gene Name** | **Sequence** |
|---|---|
| **DNMT1** | MAPTALIRQVVIDFICLAIVFVGLPLIPEKAWSTPLQGFFCNDPDIRYPYLDSSLSRILV<br>GGVGYVVVWLPVLLLHLLGKPFQRGFYCDDNSIRYPFKNSTVTNVMLYCFGLGLPIASMFI<br>VEVVRWRNNSHSKKLSRQNSCSTINLSSSVRIPSVIAEIIHLVAIFLFGAACSQLVTDFG<br>KYTIGRLRPHFIAVCQPENLAELCSSNAPHTYITNYKCTGSDEDLVRESRLSFPSGHASF<br>SAYTMLFLALYLQRRMNWSGSKLLRPTIQISALLLSWYTGLTRVSDYKHHWLLKGVLLKF<br>GSGRKHSFSEYEFLDTIMKIELVGGGEATCNLVNVKVEPVAPNPEQESSSDSSSRNTTPV<br>EEEAKPVLEQVKTKTRGVKVRYAFDDAAAIGEDDDFLPSGNSKKKRRTTSRHAKAELPDI<br>GGYRPSKKKRINRKDPNRCHVCRQRFDDPNLKYFPGPPADAIEEIIALTDPSLSLFTGEE<br>ECMSELDQRPILKLTQFGIYDEPGHLCHLDGGAIEANHLLYAYGYVKPVWNDNPGTDGGI<br>ATKEIGPINEWFISGFDGGEKAIMCFATSYADYYLMAASELYEPIMNELEEKIFLSKTVI<br>ELLIEEDNAEYEDLLTKLQNSLMPNGTRVSEDSLLRYAEFVCDRVYHFDAAGAEDEPPLI<br>LSPCMRTLIQLSGITLGKRRATRKLEKRRQPKAKKGPNWTKATTTPLVGYVFESFFRDQM<br>DQGEDKFGPTSAPRRTRCGICEACQQTDCGKCNFCRDMVKFGGSGRSKQSCALRKCPNRA<br>VQVADDDDELDAVLDSEIEVVSLDVDHKKPVKRHAYDIEWDGPELKVVDGLTFYSAAIVN<br>GDMRVFSGGHVTIEPDDPSIPMYIAEVVALWEDGKSGEQFLHARWFCRGTDTVLGETSDD<br>PRELVLIEDCEDLLLSAVVKVVNVKYKQPDPIKWKAEGGSDDPSLFQTEDDHSDTTFWYR<br>YLYHGRTGRFEDPPECPDVVNNNKGCYCCDRLDRIRQRDCAKLGNKLDSGGFDSVAWHEM<br>DIKVGDAVFLEPGAYVMRGPDGLVVKKEKIDPEEEEGFGDDYDEEYYPEKYRKTDNIKGS |

| | |
|---|---|
| | NNDTPDPFCIGYVVGVVYNGIIHNNLNAREVCLKVKRIYRPADTHLGRDAGFRSDWNLVY<br>WSDEIHNMELSKVVDKCVLVCSTAIDEPIEEFVCSGPNRMYFNKAYNPAEREFEPPPVEA<br>ERIGSSSKGKGGKSLKSAKTIQPLYPSYPKIEPLKTLDIFAGCGGLSEGLHQSGVAKTYW<br>AIESEPTAAQAFRLNNPDAAVFTDDCNTILKMAIDGHLEQNGQLLPPKNGVELLCGGPPC<br>QGFSGMNRFNSRQYSSFRNSLIVSYLSYCDYYRPRFFILENVRNFVSFKRNMVLKLTMRC<br>LVRMGYQCTFGVLQAGNYGVSQTRRRAFILAAAPGEKLPLYPEPTHVFSRRGCQLSVAVG<br>RDKFYSNCRWLLSAPYRTVTVRDAMSDLPEIPNGAKQEEISYGGDPQSHFQRWMRGTDSE<br>SSGVLRDHICKDMAPLVEARIAFIPSKPGSDWRDLPNTEVRLKDGVSTVKLRYTHEDKNG<br>RSSSGAMRGVCSCAESRQCDPLDKQHNTLIPWCLPHTGNRHNNWAGLYGRLEWDGFFSTT<br>ITNPEPMGKQGRVLHPEQHRVVSVRECARSQGFPDSYRFFGNITDKHRQVNTGYMVGNAV<br>PPPLARAIGLEIRKCVFETDSKKKKEPLTVVEI |
| **DNMT3.1** | MANKTSRSESQATENGVEPKKTLASRKFIRLGTMVWAKLDGWPWWPGIVVTHNDCGLPPP<br>RKPTNYWVYWFGGHQTVSEMPAEKLSGFLDDHFIRLLNQAHNNLYEKGVLEALRMLAHSK<br>NQPKLVKPTAGKLKSWAIKIFSSVKGNTNRDVVLDDFPEWLVRCISRIKDTNDKLKRISN<br>EKKESRLSAFQGGQKKKEKKKNNAEDDDVTYGPRDCDIAVVDRIKAGTLDMDYFCLGCHV<br>EIMGPCTEHPLFLGSLCNQECKNELLGTSYSVGEDDIHVGCAICGSTGDCLVCENGSCSR<br>VYCSYCVELLVGEGSMSAIMQKTPWFCFLCAHFATESHGLLQPRSDWALKLRNFLHQEFP<br>LQLNRHLSKNGQPSLRVVVLHDNLGAVNYALGSLRLHLDQYMASDRWEEVYELNGVDFVV<br>LKNANQMKDEEWAKLCPIHLLVSCLPAKKFDPQRHQPAGREVGRRQFAEGEYGEAFFDFF<br>HIKNCIERNNQETPLLWAVENVSCYHITISRFLQMEPIIFDIRDGQGCVKHRLVWTNIPE<br>QNFKPQLEQVINNDCPSISSLPLSREPFFIKYPWILEDIVAIAQPRPANWQEKHDRMLFQ<br>RDTRSKTKIIKRPSYLEHYSVVQLTQKSAKKRRKMDDEFHPNWRLRENGNKAEDTDDDQT<br>VYVAALTNHATYARSLGFPEPFPEFVSHLDEAAVQERLNRSLPVSLWIHLFQPLLKLARI<br>AEPDCKS |
| **DNMT3.2** | MFLSLLLSRSFSLSKMTTYDREDFYDSDATQFSESDDDNFEGNFGVDPIDDLDQWIPPLF<br>YLAKGSLQCIQNNKSSLENLCLACWEKGMHPHPFFNGGLCHPCKERLLHTMFSKSADGFH<br>LYCTVCGSRSNACVNCTSSNCIKKYCVNCLNIWTDYSDKLVNNKSNDWICFLCLPEPNLL<br>LQANHDWAQKVLFFHEPLSVYGPRALYWKKQPLRVLSLFDGIGTGLVALRKLGIQVEVYY<br>ASEVLTAAATVSRTRLGGVLQHIGSVGEVTERRLEEIAPIHLLIGGSPCNDFSAINRFPK<br>DFYDPRGYSRYFFDFVRVLNLMRKINGQYQHLLWLFENVASMPQHYRDTISRHLDCQPAV<br>IDAKNFSPQLRRRLFWGNIPGLFTVHAEQMTMDGELLTLDKSLMPNSGRSAAQAKIRTLT<br>TNTNSLLQGRTENCKSRKDLASLFPVRFSFQQDELRNADVENDVARHSKKGKRDSTSGIN<br>ATDKPPRTVADEDDNQQTDVLWLQEIEHVFGLPRHFTDVGNMSRTDRQKLLGHAWSVPVI<br>VSIFSNLKSYTV |
| **UHRF1** | MDGKDSVMLTVSKMSLIEEVRQLVKDKLNVDPVCQRLFFRGKQMEDGYRLIDYGININDV<br>VQLMVRAIPVPQPIKEVNEEGSEIEETTVDKKEAKKSGDESLTDAICEYYEVQDLIDAKD<br>PFTSSWVEAKIVRIAKNKEDDPLEYHVLFQGHEREIPLPRSFQQIRPRAEELCSNADLKL<br>GEVVLVNYNMEESKERGLWYDGKLTKADLQSRTKKKVIVTLIMGEENETEVNDCSIYFVK<br>EIMRIPQVKKRRDQTAEETRLLKKGPATKRESAPYCHHCNDNPRRKCKFCGCHECGSKEN<br>PDQQIMCDECDLPYHLYCLKPPLTCMPDESEEWYCPKCKTDSSQIVKAGEKLKESKKKAK |

| | |
|---|---|
| | APSNLNKNTTRDWGRGMACAGRAKECTIVPSNHFGPIPGVDVGTTWRFRFQASEAGVHRP<br>PVGGIHGREKEGAYSIVLSGGYEDDMDNGDSFYYTGSGGRDLTGNKRTAGQSCDQTLTRM<br>NLALALNCNVEVNETNGAEAKDWRKGKPVRVLRKGHADEKSLAKGPSKGKGKAAKHASSY<br>GPEIGVRYDGIYKIVKYWPEKGKSGFLVWRYFLQRDDPTPPVWTEEGKKRIQQLGLDHVI<br>YPEGHLEAMEAKEQEKEKNGGKRKSVVELLQQNKESSTAKKAKKVGYQLEPEIAELIEKD<br>ILNRKLWDECKESLDDTKHKFVSKVEERFLCICCQEIVCKPITTSCTHNICLACLQGSFR<br>AKVFTCPSCRHELGKNMAMEPNENLCRALNAIFPGYENGR |

**Table S2. DNA extraction and alignment results of 2 deeply sequenced *D. magna* samples and 15 *D. magna* samples.** Dap_S1 and Dap_S2 are two deeply sequenced samples and are categorized as the mature-old group. Dap_D1 to Dap_D4 are categorized as the young group; Dap_D5 to Dap_D8 and Dap_D13 to Dap_D16 are categorized as the mature group (Dap_D7 was excluded due to low quality); and Dap_D9 to Dap_D12 are categorized as the old group.

| Sample | Age (day) | Fastq Reads Paired | Alignment after Filtering | Alignment after Filtering (%) |
|---|---|---|---|---|
| Dap_S1 | 45 | 584,073,458 | 300,104,273 | 0.51381255 |
| Dap_S2 | 45 | 567,906,032 | 312,642,898 | 0.550518713 |
| Dap_D1 | 9 | 39,844,366 | 24,415,816 | 0.612779634 |
| Dap_D2 | 9 | 47,652,374 | 25,427,798 | 0.533610309 |
| Dap_D3 | 9 | 52,740,124 | 29,991,863 | 0.568672592 |
| Dap_D4 | 9 | 42,613,178 | 25,005,893 | 0.586811268 |
| Dap_D5 | 25-27 | 39,513,448 | 24,172,883 | 0.611763443 |
| Dap_D6 | 25-27 | 35,855,664 | 21,647,932 | 0.603752088 |
| Dap_D8 | 25-27 | 35,907,888 | 22,704,957 | 0.632311123 |
| Dap_D9 | 58 | 45,600,500 | 27,883,304 | 0.61146926 |
| Dap_D10 | 58 | 25,608,764 | 14,629,599 | 0.571273139 |
| Dap_D11 | 51-52 | 42,469,308 | 26,206,058 | 0.617058747 |
| Dap_D12 | 51-52 | 59,805,218 | 35,731,844 | 0.597470341 |
| Dap_D13 | 22-23 | 48,905,768 | 30,357,887 | 0.620742465 |
| Dap_D14 | 22-23 | 68,368,080 | 40,374,256 | 0.590542487 |
| Dap_D15 | 22-23 | 47,102,532 | 28,952,651 | 0.614672922 |
| Dap_D16 | 22-23 | 52,749,350 | 31,873,233 | 0.604239351 |

**Table S3. Coverage summary of cytosine sites in *D. magna* samples.** All cytosine sites included in downstream analysis have a minimum coverage of 10x.

| Sample | Min | 1st Qu. | Median | Mean | 3rd Qu. | 90 perc. | 99 perc. | Max |
|---|---|---|---|---|---|---|---|---|
| Dap_S1 | 10 | 48 | 65 | 67.85 | 83 | 103 | 162 | 6943 |
| Dap_S2 | 10 | 47 | 64 | 68.24 | 83 | 107 | 178 | 6749 |
| Dap_D1 | 10 | 10 | 11 | 12.78 | 13 | 16 | 30 | 3365 |
| Dap_D2 | 10 | 10 | 12 | 13.40 | 14 | 17 | 33 | 4648 |
| Dap_D3 | 10 | 11 | 12 | 13.29 | 14 | 17 | 31 | 4213 |
| Dap_D4 | 10 | 10 | 12 | 13.23 | 14 | 17 | 32 | 3692 |
| Dap_D5 | 10 | 10 | 11 | 12.95 | 13 | 16 | 32 | 3535 |
| Dap_D6 | 10 | 10 | 11 | 12.93 | 13 | 16 | 33 | 3220 |
| Dap_D8 | 10 | 10 | 11 | 12.82 | 13 | 16 | 31 | 2679 |
| Dap_D9 | 10 | 10 | 12 | 13.10 | 14 | 17 | 31 | 3357 |
| Dap_D10 | 10 | 10 | 11 | 13.43 | 13 | 16 | 38 | 1986 |
| Dap_D11 | 10 | 10 | 12 | 13.04 | 14 | 17 | 30 | 2936 |
| Dap_D12 | 10 | 11 | 13 | 13.86 | 15 | 18 | 32 | 3294 |
| Dap_D13 | 10 | 11 | 12 | 13.34 | 14 | 17 | 30 | 3533 |
| Dap_D14 | 10 | 11 | 13 | 14.52 | 16 | 19 | 33 | 4219 |
| Dap_D15 | 10 | 11 | 12 | 13.27 | 14 | 17 | 31 | 3344 |
| Dap_D16 | 10 | 11 | 12 | 13.42 | 14 | 17 | 30 | 4276 |

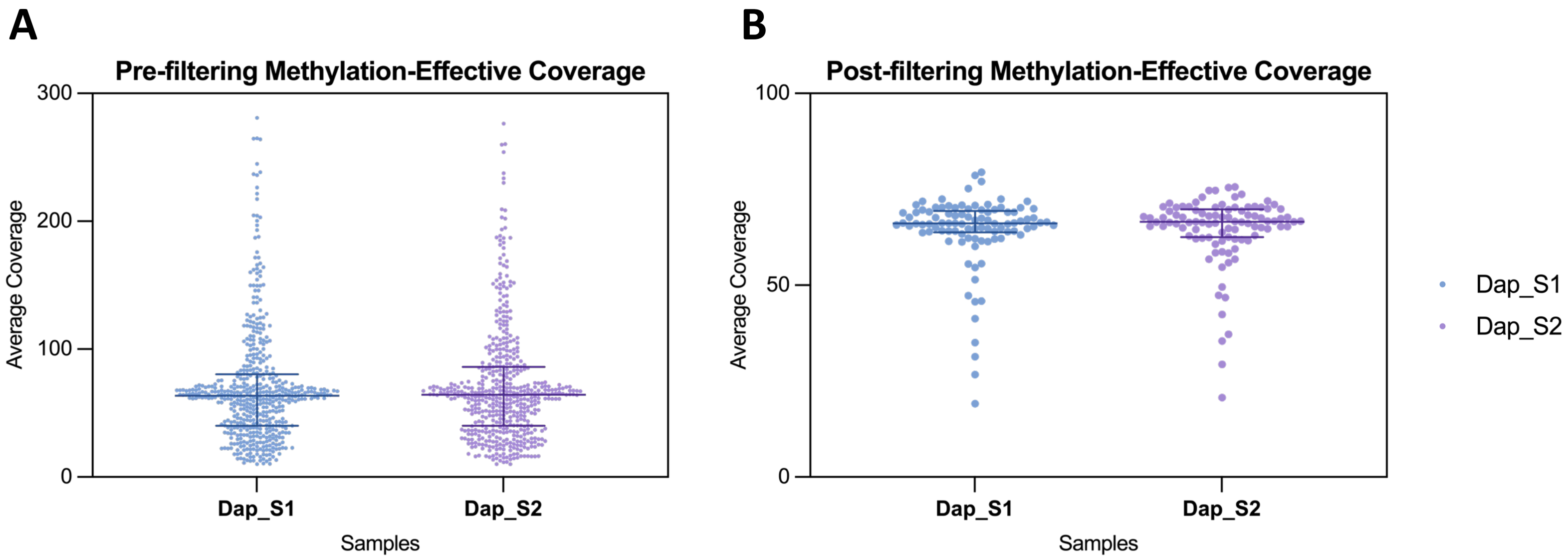


**Figure S1. Pre-filtering and post-filtering distribution of average methylation-effective coverages per contig in two deeply sequenced samples.** The methylation-effective coverages per contig were calculated as the average read coverage across all CpN sites on a contig. **(A)** Before filtering, the 608 nuclear contigs had a minimum average coverage of 10 and an approximate median average coverage of 60 and demonstrated with a variety in coverage values of contigs. **(B)** The average coverage of 97 contigs after filtering centered around 60 with less variability and consistency across samples.

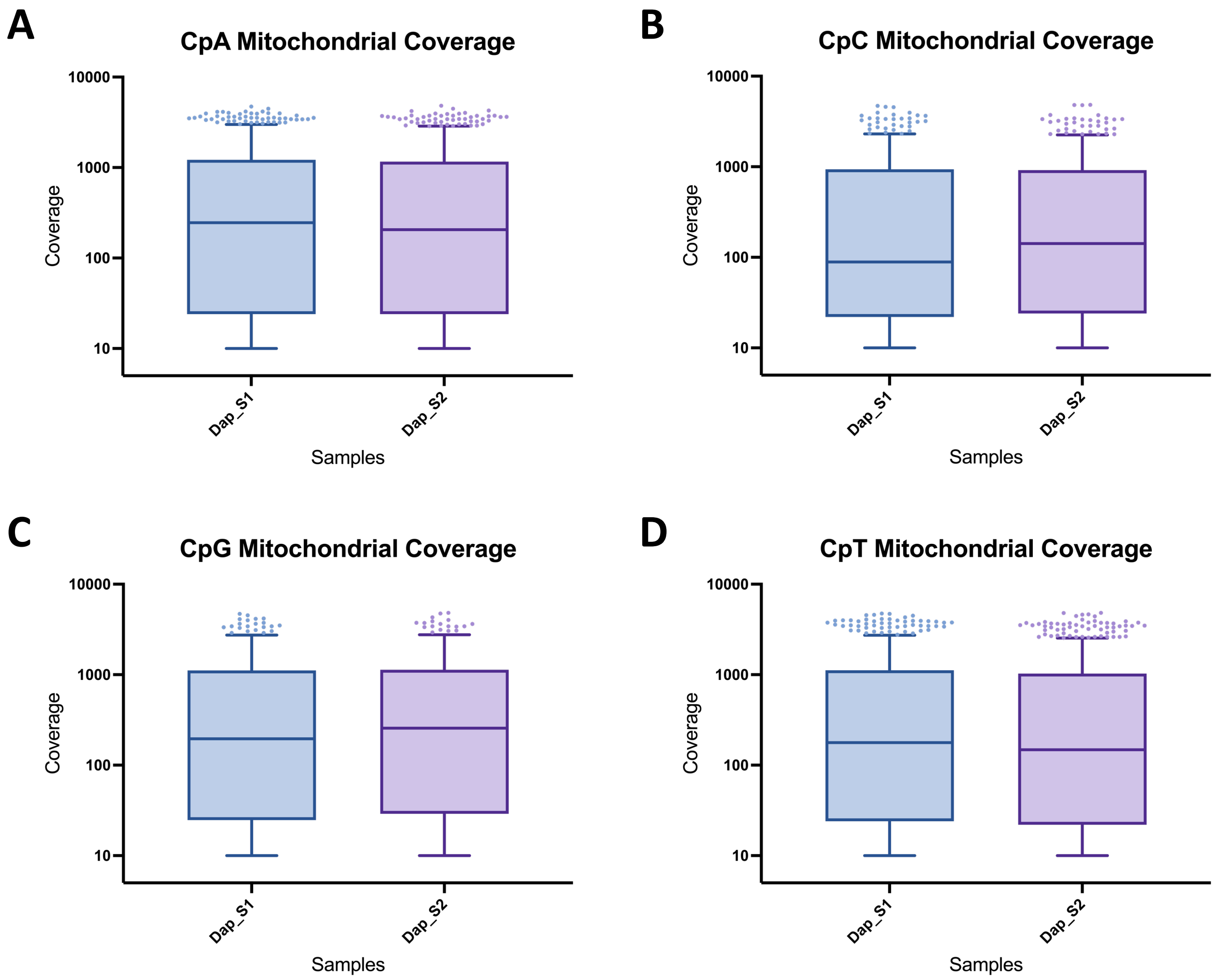


**Figure S2. Mitochondrial genome coverage for two deeply sequenced samples.** Boxplots show the distribution of methylation coverage for four cytosine contexts, **(A)** CpA, **(B)** CpC, **(C)** CpG, and **(D)** CpT, in the mitochondrial genome of Dap_S1 and Dap_S2. Methylation coverage is generally comparable between the two samples across all cytosine contexts.

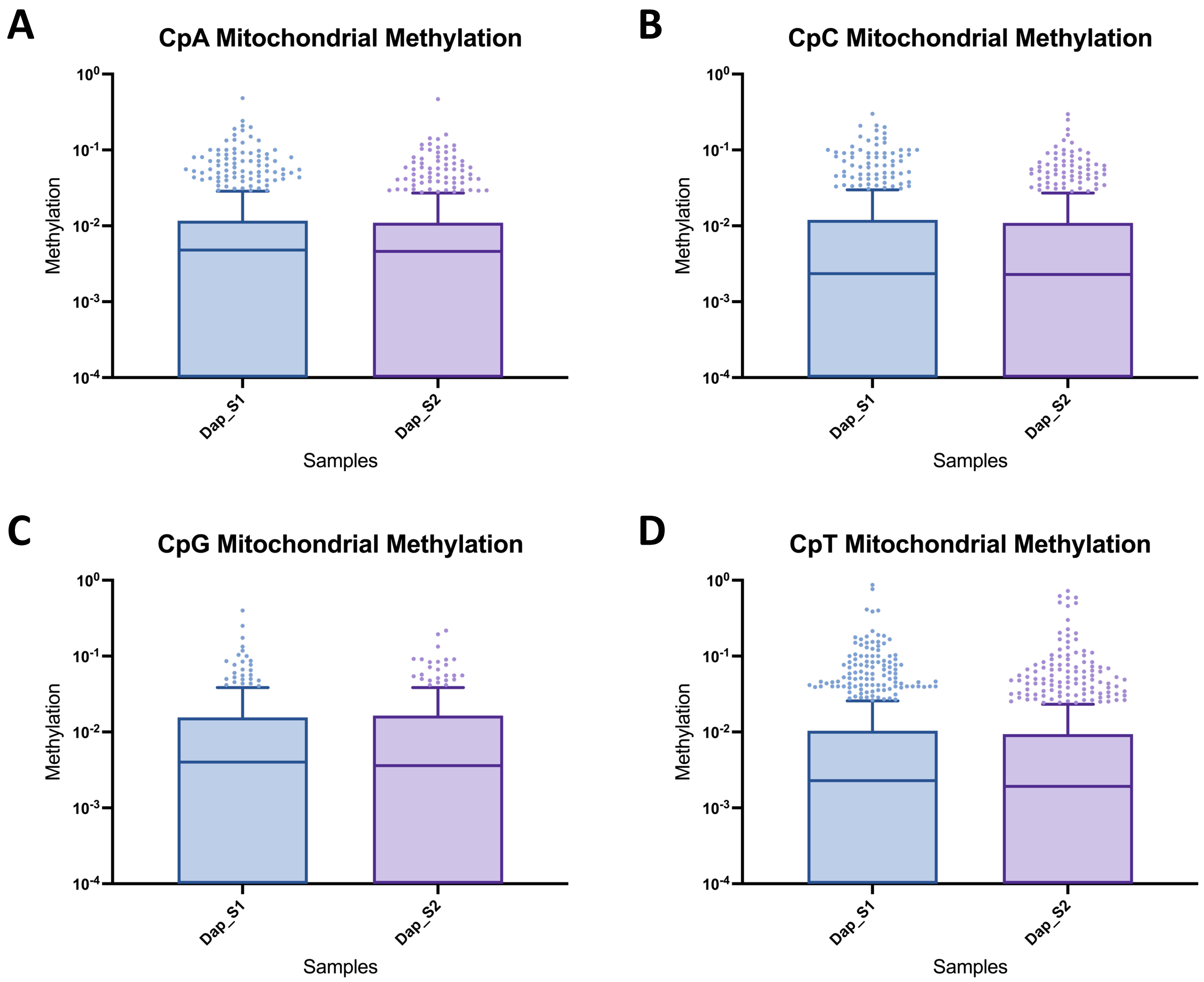


**Figure S3. Mitochondrial genome cytosine methylation levels for two deeply sequenced samples.** Boxplots illustrate the distribution of methylation levels across four cytosine contexts, **(A)** CpA, **(B)** CpC, **(C)** CpG, and **(D)** CpT, in the mitochondrial genome of Dap_S1 and Dap_S2. Methylation levels are generally consistent between the two samples across all cytosine contexts. CpG sites exhibit the fewest outliers with high methylation levels, suggesting a more uniform distribution in comparison to the other cytosine contexts.

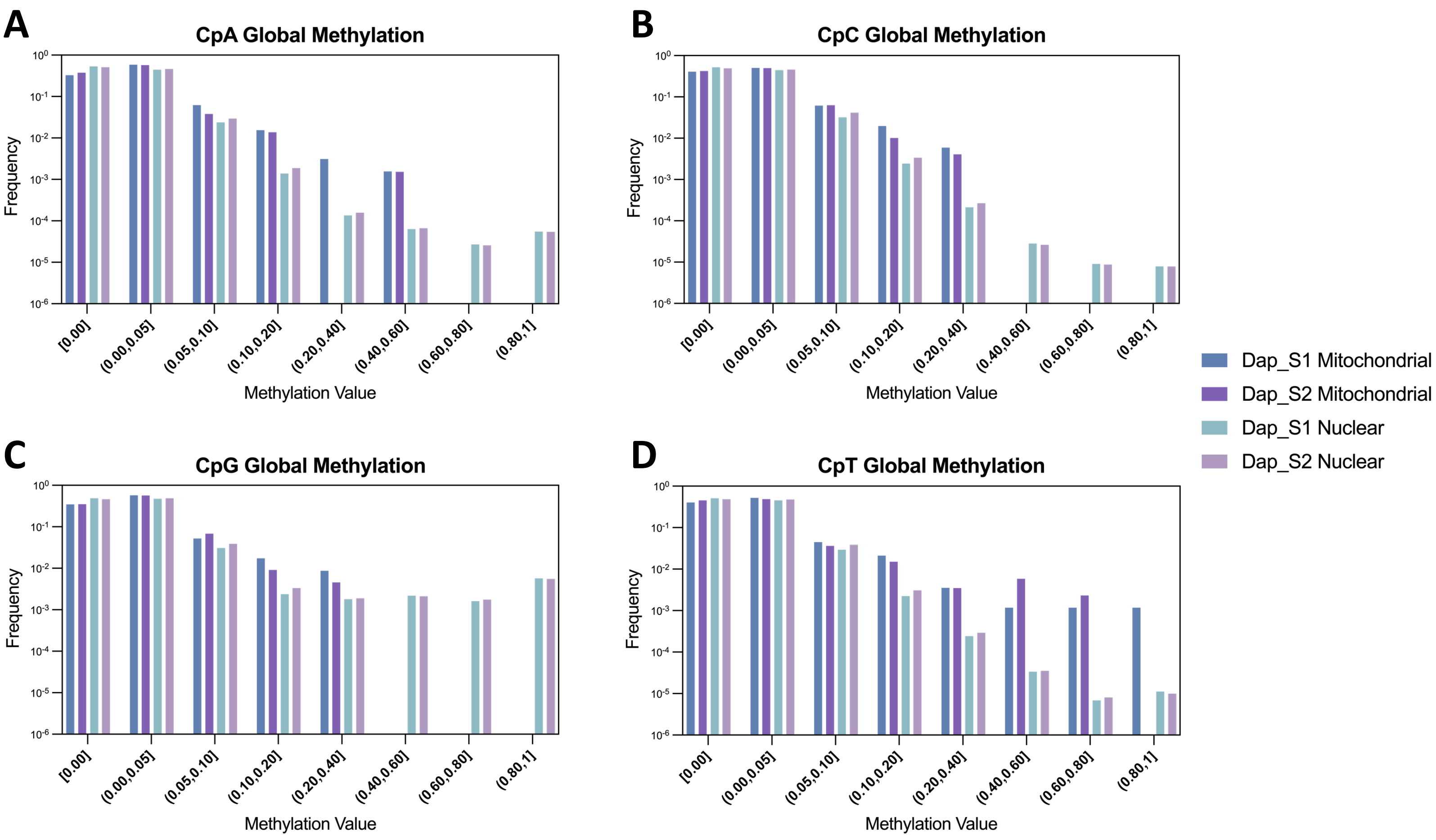


**Figure S4. Mitochondrial and nuclear genomic cytosine methylation levels for two deeply sequenced samples.** Methylation levels of individual CpN sites from mitochondrial or nuclear genome are presented as the frequency distribution. **(A)** CpA, **(B)** CpC, and **(D)** CpT sites in mitochondria exhibit higher average methylation, while **(C)** CpG sites exhibit a significant higher degree of nuclear methylation.

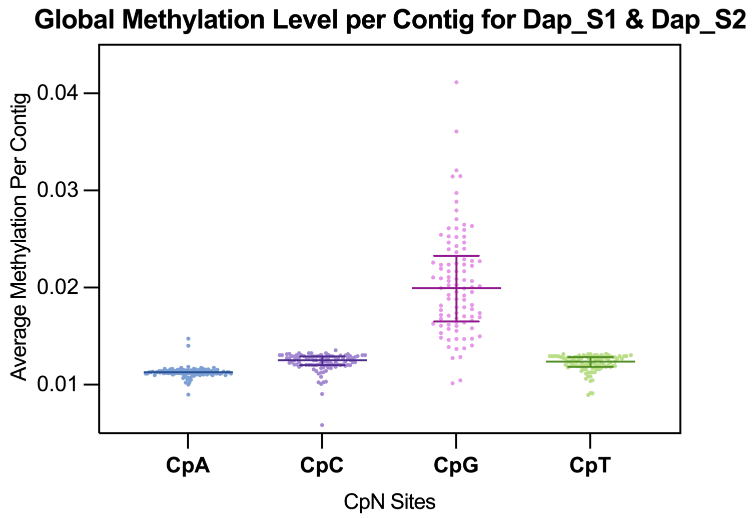


**Figure S5. Global methylation levels per contig for two deeply sequenced samples.** Scatter plot show the average methylation levels per contig for four cytosine contexts: CpA, CpC, CpG, and CpT, in both Dap_S1 and Dap_S2 samples. CpG sites displayed higher average methylation levels across contigs, whose distribution of methylation was significantly different from that of the non-CpG sites.

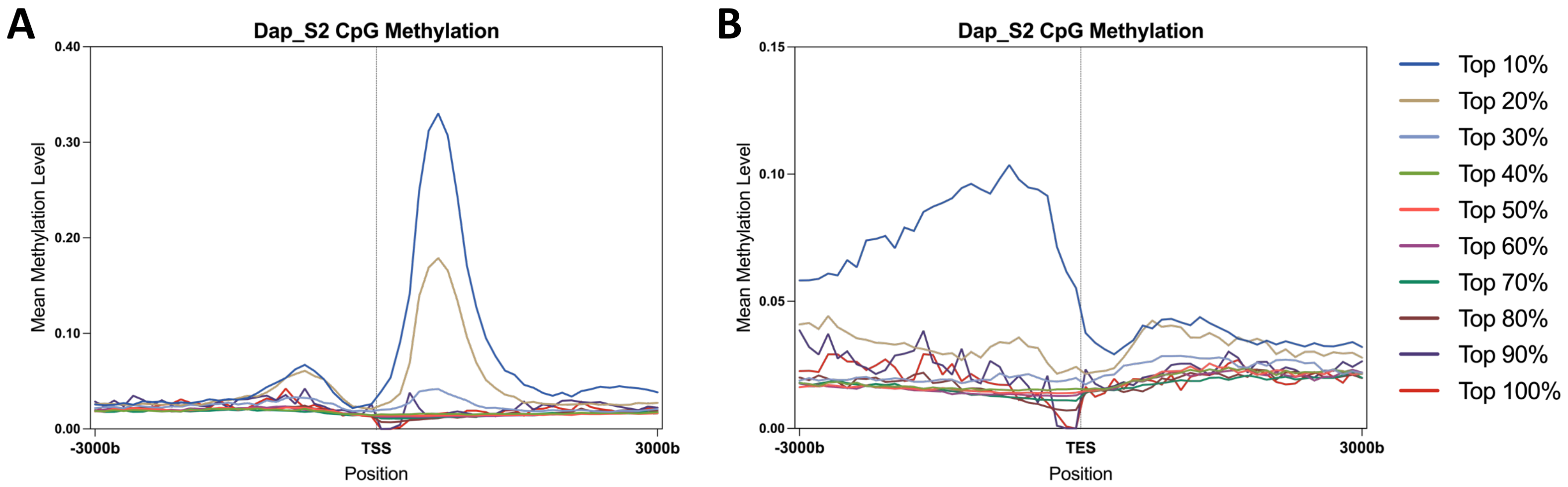


**Figure S6. Gene-level CpG Methylation Patterns Centered around TSS and TES for Dap_S2, spanning 3 kilobases (kb) upstream and downstream of transcription start sites (TSS) and transcription end sites (TES).** Genes were divided into 10 groups based on their mean methylation levels. Each line represents the average methylation level within each group, centered around **(A)** the transcription start site (TSS) and **(B)** the transcription end site (TES). For example, Top 20% in legend represents the top 10-20% group.

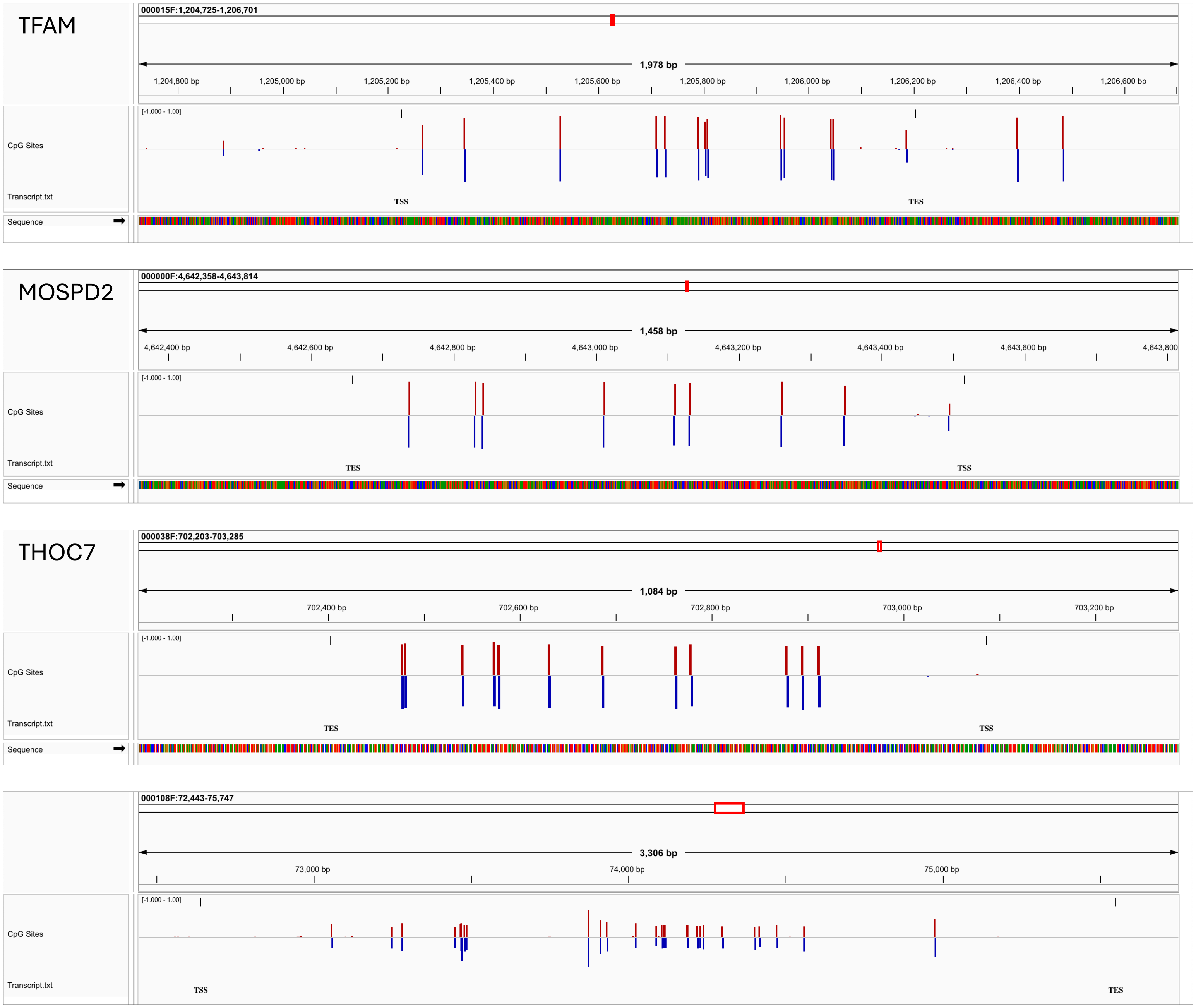


**Figure S7. Genome Browser Snapshots of Genes Top 10% Methylated in Dap_S1.** The genome browser snapshots display full genes and nearby regions, offering a view of CpG site positions and methylation levels. CpG sites on the Watson (sense) strand are represented by red bars, while those on the Crick (antisense) strand are shown with blue bars. The gene labels correspond to human homolog mapped to *D. magna* sequences.

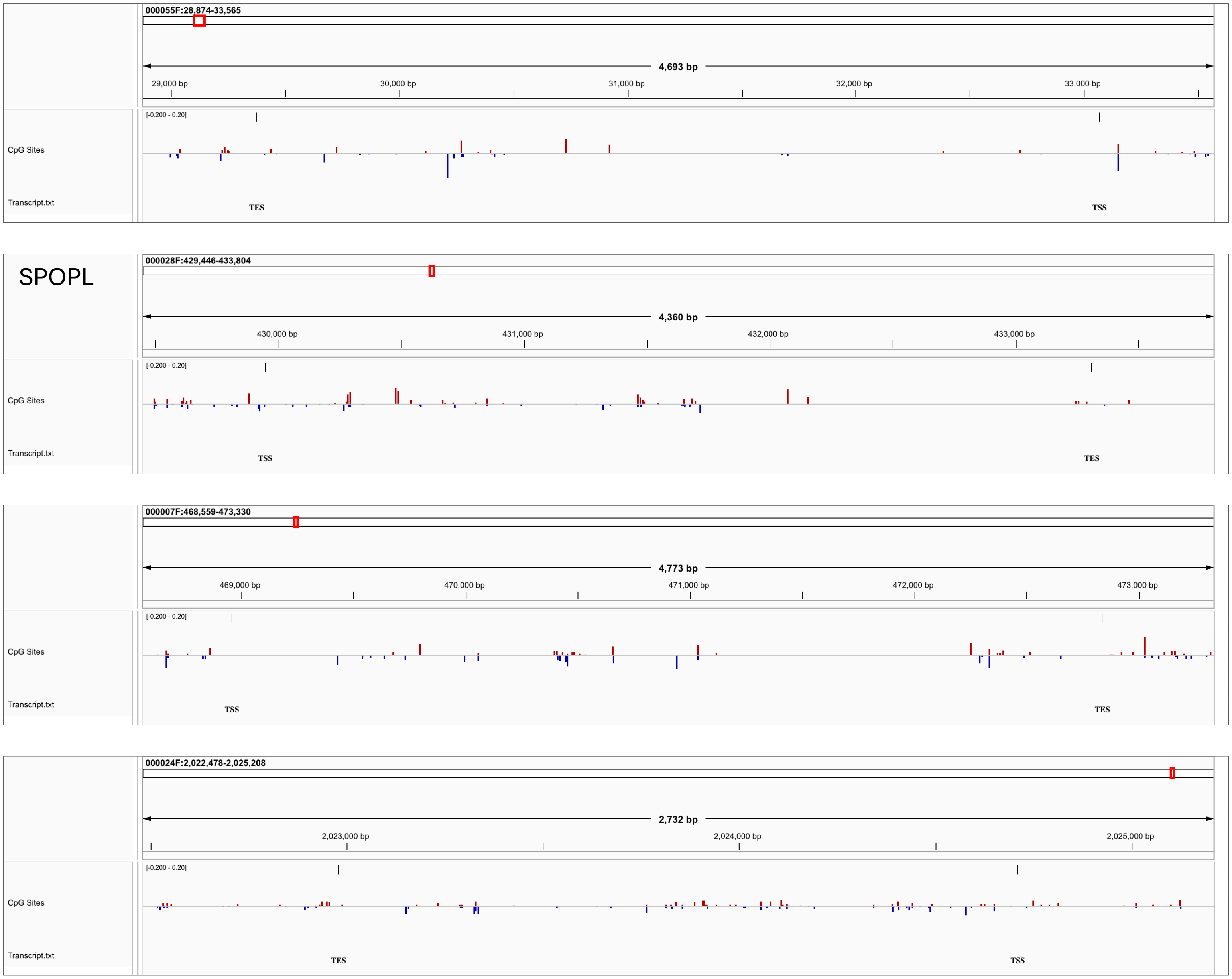


**Figure S8. Genome Browser Snapshots of Genes Lower 50% Methylated in Dap_S1.** The genome browser snapshots display full genes and nearby regions, offering a view of CpG site positions and methylation levels. CpG sites on the Watson (sense) strand are represented by red bars, while those on the Crick (antisense) strand are shown with blue bars. The gene labels correspond to human homolog mapped to *D. magna* sequences.

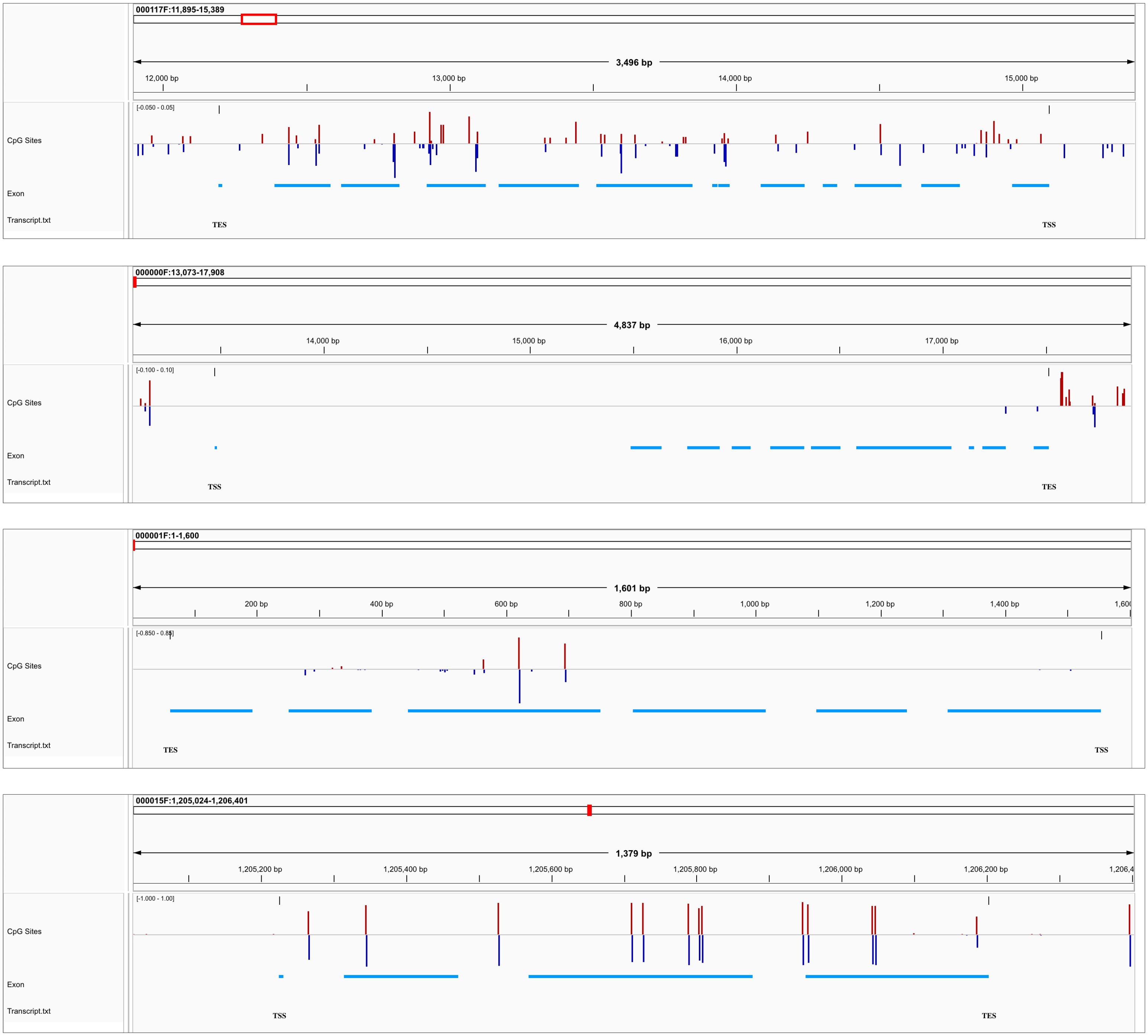


**Figure S9. Genome Browser Snapshots of Genes and Exons in Dap_S1.** The genome browser snapshots display full genes and their exons (light blue bars), offering a view of CpG site positions and methylation levels. CpG sites on the Watson (sense) strand are represented by red bars, while those on the Crick (antisense) strand are shown with blue bars. Exons are more methylated for CpG sites compared to introns.

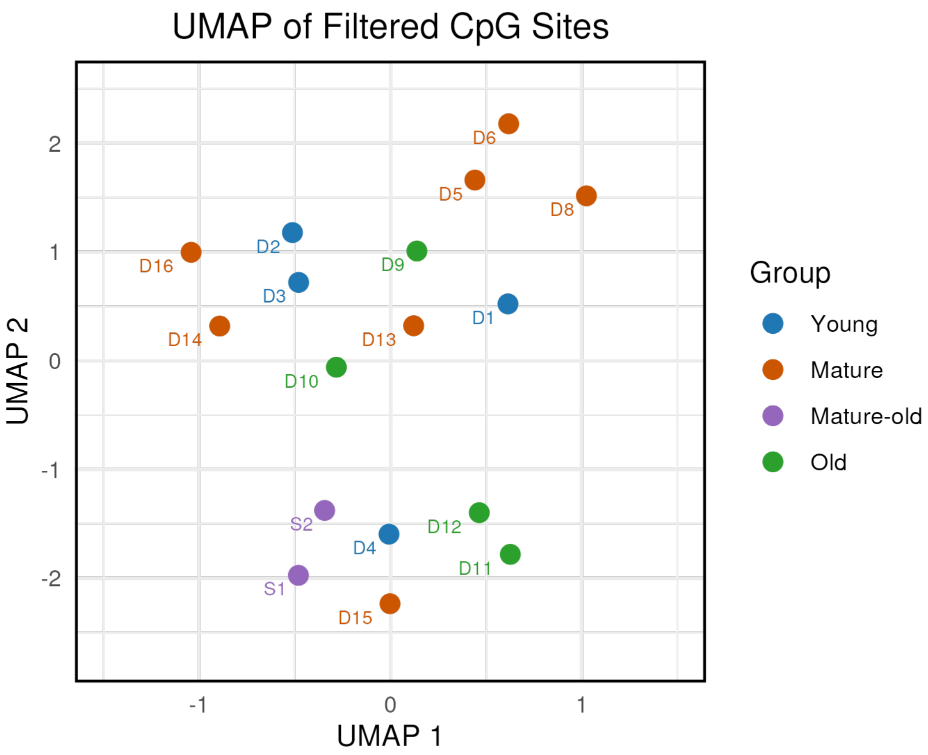


**Figure S10. CpG methylation from selected genes demonstrates no age-related clustering.** CpG sites present in genes with at least 10% methylation in at least 80% of the 17 samples were retained. The UMAP visualization of matrix with selected CpG sites does not reveal any specific age-related clustering.

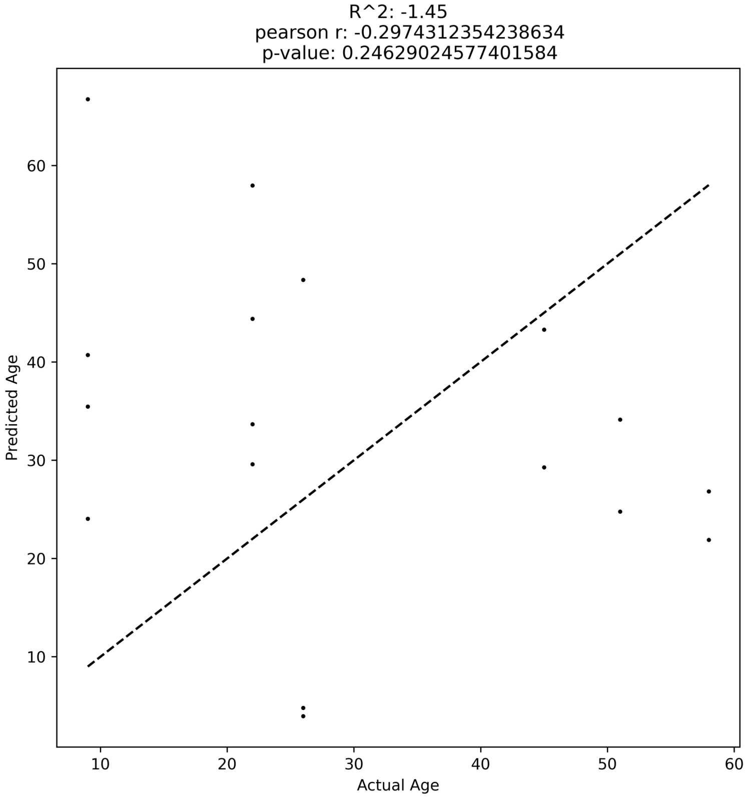


**Figure S11. Epigenetic clock model constructed using a CpG methylation matrix.** CpG sites present in at least 80% of the 17 samples were included in the matrix, and missing values were imputed. The CpG matrix was filtered to retain the top 20% of the CpG sites with the highest variability. A Lasso regression model was then applied to predict age based on CpG methylation levels. The resulting epigenetic clock showed a low and statistically insignificant correlation between predicted age and actual age.

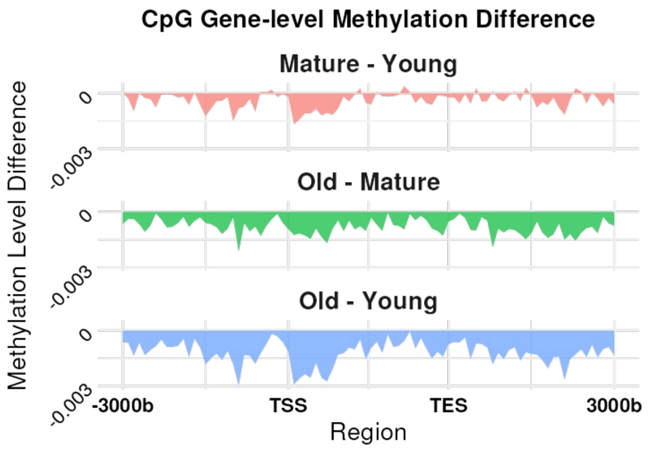


**Figure S12. Gene-level CpG methylation patterns reveal subtle decreases with age.** Differential CpG methylation levels between age groups show a weak trend of decreasing methylation with advancing age across gene bodies and neighboring regions.
